# Mechanical Coupling Coordinates the Co-elongation of Axial and Paraxial Tissues in Avian Embryos

**DOI:** 10.1101/412866

**Authors:** Fengzhu Xiong, Wenzhe Ma, Bertrand Bénazéraf, L. Mahadevan, Olivier Pourquié

## Abstract

Tissues undergoing morphogenesis impose mechanical effects on one another. How developmental programs adapt to or take advantage of these effects remains poorly explored. Here, using a combination of live imaging, modeling, and microsurgical perturbations, we show that the axial and paraxial tissues in the forming avian embryonic body coordinate their rates of elongation through mechanical interactions. First, a cell motility gradient drives paraxial presomitic mesoderm (PSM) expansion, resulting in compression of the axial neural tube and notochord; second, elongation of axial tissues driven by PSM compression and polarized cell intercalation pushes the caudal progenitor domain posteriorly; finally, the axial push drives progenitors to emigrate into the PSM to maintain tissue growth and cell motility. These interactions form an engine-like positive feedback loop, which ensures the tissue-coupling and self-sustaining characteristics of body elongation. Our results suggest a general role of inter-tissue forces in the coordination of complex morphogenesis involving distinct tissues.

## INTRODUCTION

During development, the morphogenesis of one tissue can produce forces that directly impact the dynamics of its neighboring tissues (Savin et al., 2011; Lye et al., 2015; Morita et al., 2017). These interactions may play a role not only in controlling the correct development of each tissue and organ but also in coordinating them into an integrated system. Such role of inter-tissue forces is poorly explored due to the challenge of experimentally detecting and applying such forces and the lack of a framework that bridges developmental signals, cell dynamics and tissue mechanics.

Avian embryos are an ideal vertebrate system to address these challenges because of their large size and accessibility. The forming body axis provides a striking example of coordinated morphogenesis of multiple distinct tissues (Bénazéraf and Pourquié, 2013). The axis forms in a head to tail (anterior to posterior, AP) sequence where tissues derived from the three germ layers are progressively added from a posterior growth zone (e.g. primitive streak, tailbud). The posterior movement of the growth zone lays down in its trail a conserved pattern of tissues which include the axial organs: neural tube (NT) and notochord (NC), and the flanking paraxial tissues: presomitic mesoderm (PSM), intermediate mesoderm and lateral plates (Figures 1A-B). While these tissues achieve similar lengths at the same pace, they exhibit very distinct cellular organization and elongation dynamics (Keller and Danilchik, 1988; Shih and Keller, 1992; Bénazéraf and Pourquié, 2013; Steventon et al., 2016; Colas and Schoenwolf, 2001; Glickman et al., 2003; Sausedo and Schoenwolf, 1993; Yang, et al. 2002; Bénazéraf et al., 2010; Dray et al., 2013; Bénazéraf et al., 2017). First, the elongation of the NC and NT largely relies on intrinsic cell polarity-driven medio-lateral intercalation promoting convergence and extension along the AP axis. In contrast, the adjacent PSM shows limited convergence in the posterior body axis region (Shih and Keller, 1992; Bénazéraf and Pourquié, 2013). Second, in NC and NT, cells exhibit an epithelial organization, move coherently and only limited cell rearrangements are seen, while in the PSM cells are mesenchymal, show random cell movements and extensive cell mixing is observed (Colas and Schoenwolf, 2001; Bénazéraf et al., 2010). Third, both NT and PSM rely on new cell addition from a specific subdomain of the growth zone called the chordo-neural hinge (or PD for progenitor domain) at the posterior end of the axis (Figure 1A). To enter the PSM, progenitors move laterally away from the medially located PD (Yang et al., 2002). In contrast, NC lengthens mainly through cell rearrangement and differentiation with little addition of new cells (Sausedo and Schoenwolf, 1993; Bénazéraf et al., 2017). These differences in cell behaviors lead to counterintuitive cellular dynamics such as sliding between PSM and NC cells with NC cells moving faster posteriorly than PSM cells, despite the two tissues appearing to elongate hand-in-hand (Figure 1B, Yang et al., 2002; Glickman et al., 2003; Bénazéraf et al., 2017). These observations raise the question of how do the PSM, NC and NT, which employ such distinct cellular processes to lengthen, coordinate their elongation during body axis formation?

**Figure 1.**
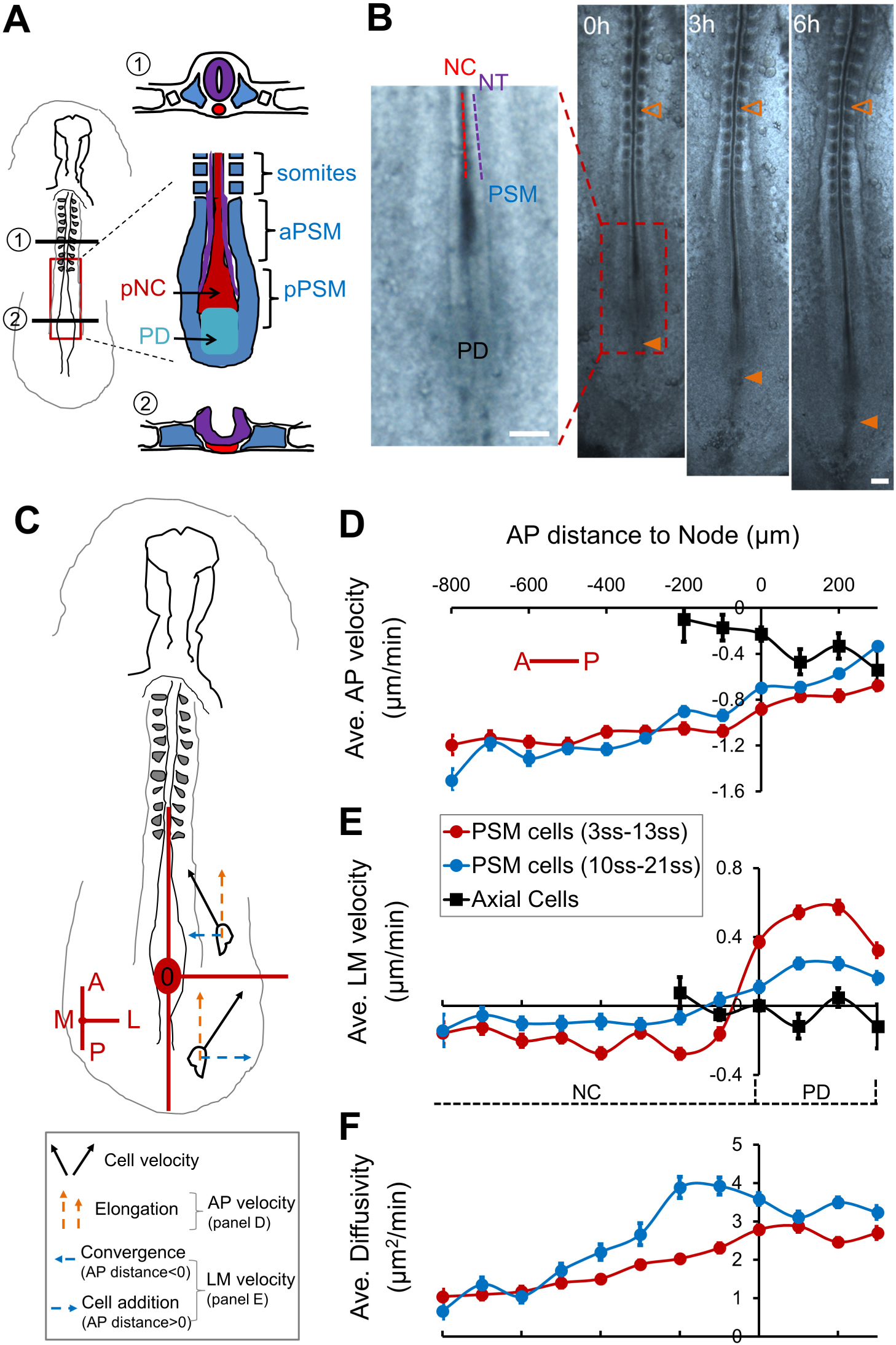
Cellular dynamics of tissues in the elongation avian body axis. (A) Diagrams of tissues of interest in the context of the whole embryo (left, anterior to the top), showing a ventral view (right) and cross-sections of formed (①) and forming (②) body axis, respectively. PD, progenitor domain; NC, notochord; PSM, presomitic mesoderm; In ①, the uncolored small squares by the somites represent the intermediate mesoderm, the further lateral uncolored blocks represent the lateral plate mesoderm. A lowercase letter “p”(“a”) in front of NC/NT/PSM indicates “posterior” (“anterior”), same in other figures. (B) A developing embryo (ventral view, anterior to the top, same as in [A]) showing axis elongation. Box: tissue domains of the axis end where new axis forms. Empty arrowheads mark the same somite as a reference; filled arrowheads marks the PD. Note the distance increase between arrowheads (elongation). Scale bars: 200μm. (C) Schematic view of the reference axes used for representing cell dynamics data. The Node area is set as the origin (0,0). A, anterior; P, posterior; M, medial; L, lateral. In this reference elongation is represented by posterior to anterior movement of PSM cells as they are left behind by the node (orange arrows). The PD level posterior to the Node has a positive AP position while the NC level anterior to the Node has a negative AP position. (D-E) Average cell speed along the AP (D) and LM axes (E). Along AP, negative indicates P to A movement in the perspective of the Node (0,0). Along LM, negative and positive indicate convergent (L to M) and dispersive (M to L) movements, respectively. Each bin integrates 100μm along the AP axis averaging data points from all time points of the movie. Error bars: ±s.e.m. (F) Cell “Diffusivity” obtained by fitting the mean square displacement (MSD) of single cell tracks with a drifting speed (directional) and a diffusion component (random).

Mechanical interactions between tissues might offer an answer to this question. The close proximity between the PSM, NC, NT and the PD means that the deformation of one would likely have a direct mechanical effect on the others. Indeed, it was found that ablating the posterior PSM (pPSM) greatly reduces the elongation rate of all tissues (Bénazéraf et al., 2010). This finding suggests that the pPSM may produce forces that contribute to axial elongation. It was further hypothesized that cell motility in the PSM underlies force generation and posterior tissue expansion (Regev et al., 2017). To test these possibilities, tissue and cell behaviors need to be observed *in vivo* with readout of tissue forces.

Here we used soft gels to replace different tissues in the elongating posterior body axis of chicken embryos to track the forces applied by the neighboring tissues. We found that the pPSM compresses the axial tissues. This extrinsic force adds to the intrinsic process of cell intercalation to promote convergence and elongation of the NT and NC. The compression requires FGF controlled cell motility in the pPSM while the NT cell intercalation requires Wnt/PCP (planar cell polarity) mediated cell polarity. Using an agent(cell)-based model that incorporates cell motility and polarity, we faithfully recapitulate the dynamics of cells and tissues under different experimental conditions. The model predicts that the advancing axial tissues create a pushing force on the PD, which we validated by gel deformation analysis. Surprisingly, the model also predicts that the cell emigration from the PD to the pPSM requires this axial pushing force. To test this hypothesis, we inserted a magnet controlled pin in place of the posterior NC/NT (pNC/pNT) region to apply a posteriorly oriented force to the PD. This mechanical signal partially restores cell emigration and posterior movement of the PD (i.e. axial elongation). Together our findings reveal mechanical coupling between the PSM and axial tissues: the left and right pPSMs generate a bilateral compression of the axial tissues promoting their elongation. The axial tissues in turn push on the PD promoting PSM growth by new cell addition. These interactions form a self-sustaining positive feedback loop that prevents elongation offset from occurring between different tissues. Just like an internal combustion engine that self-refills fuel for the next cycle, expanding pPSM (combustion) forces axial tissues (piston) to drive PSM progenitors (fuel) out of the PD (fuel reserve). Our model thus provides a simple physical mechanism and an intuitive explanation of the striking robustness and coordination between tissues observed in embryonic body axis formation.

## RESULTS

### Cellular dynamics of axial and paraxial elongation

To set up a quantitative baseline of tissue and cell behaviors during body axis elongation, we performed live imaging of Tg(CAG:GFP) chicken (McGrew et al., 2008) embryos after labeling the mesodermal progenitors in the primitive streak and tailbud with DiI (Figures S1A-B, Movie S1) between Hamburger Hamilton (HH) stages 8 and 14. During these stages, body elongation first follows the regression of the primitive streak and then the posterior movement of the tailbud. These movements result in the progressive formation of the NT, NC and PSM (among other tissues) at the posterior end of the embryo (Hamburger 1992).

Using cell tracking (Figure S1C), we analyzed the movement of cells relative to the Node (Figure 1C). Both axially-localized cells (including NC cells and PSM progenitors in the PD) and paraxially-localized PSM cells are left behind by the Node but at increasingly different speeds (Figure 1D, note that while some axial cells move posteriorly with the Node, others are left behind, resulting in a net negative average speed at the Node [position 0]). This speed offset reflects the rate of the posterior movement of the Node and the PD (i.e. the elongation rate). In early embryos, at the PD level of the PSM, cells move away from the midline while in the PSM flanking the NC, cells converge toward the midline (Figure 1E, positive indicates medial to lateral movement). Similar dynamics occur in later stage embryos but at markedly reduced average speeds (Figure 1E). The medial to lateral movement of progenitors at the PD level results in their joining the PSM on both sides of the axis (Figures S1D). The average speed of this movement thus serves as an indicator of PSM cell addition rate (Figure 1E). To assess the motility of these cells (Figures S1E), we fit the cell tracks with a diffusion model to separate the directional and non-directional components (Regev et al., 2017). This allows us to assess the local motility or “cell diffusivity” in the PSM. A posterior to anterior motility gradient is observed in both early and later stages of elongation (Figure 1F). These cellular dynamics are consistent with previous studies (Kulesa and Fraser, 2002; Yang et al., 2002; Manning and Kimelman, 2015; Bénazéraf et al., 2010, 2017).

### Posterior PSM exerts a lateral to medial compression on the axial tissues

To investigate how the pPSM affects the axial tissues, we performed bilateral ablations of the pPSM (Figure 2A). Interestingly, both pNC and pNT remained wider even after PSM cuts closed, suggesting impaired tissue convergence (Figures 2B-C). Correspondingly, the elongation rate of the operated embryos reduces drastically (Figures S2A-C). These data show that the pPSM is required for the convergence and elongation of the axial tissues. To test if the pPSM functions as a barrier that passively confines pNC/pNT to prevent their lateral expansion, we replaced the pPSM with a stiff alginate gel. The gel-flanked portions of pNC/pNT remained wider (Figure 2D, n=4/4), suggesting that the pPSM actively compresses the axial tissues. To detect this compression, we replaced the pNC/pNT with a soft alginate gel (Gelation is performed using 1% alginate at 10mM Ca^2+^ or *in vivo* to reduce gel stiffness to the order of 100 Pa [Banerjee et al., 2009] comparable to embryonic tissues [Zhou et al., 2009]). Using live confocal imaging (Figure 2E, Movies S2), we found that the gel narrows significantly over time along the LM axis (Figures 2E-G). Similar deformation dynamics are observed in replacement experiments of the pNC (Figures S2D-G) or the pNT (Figure S2H) alone. These results show that pPSM contributes to axial convergence and elongation through lateral tissue compression.

**Figure 2.**
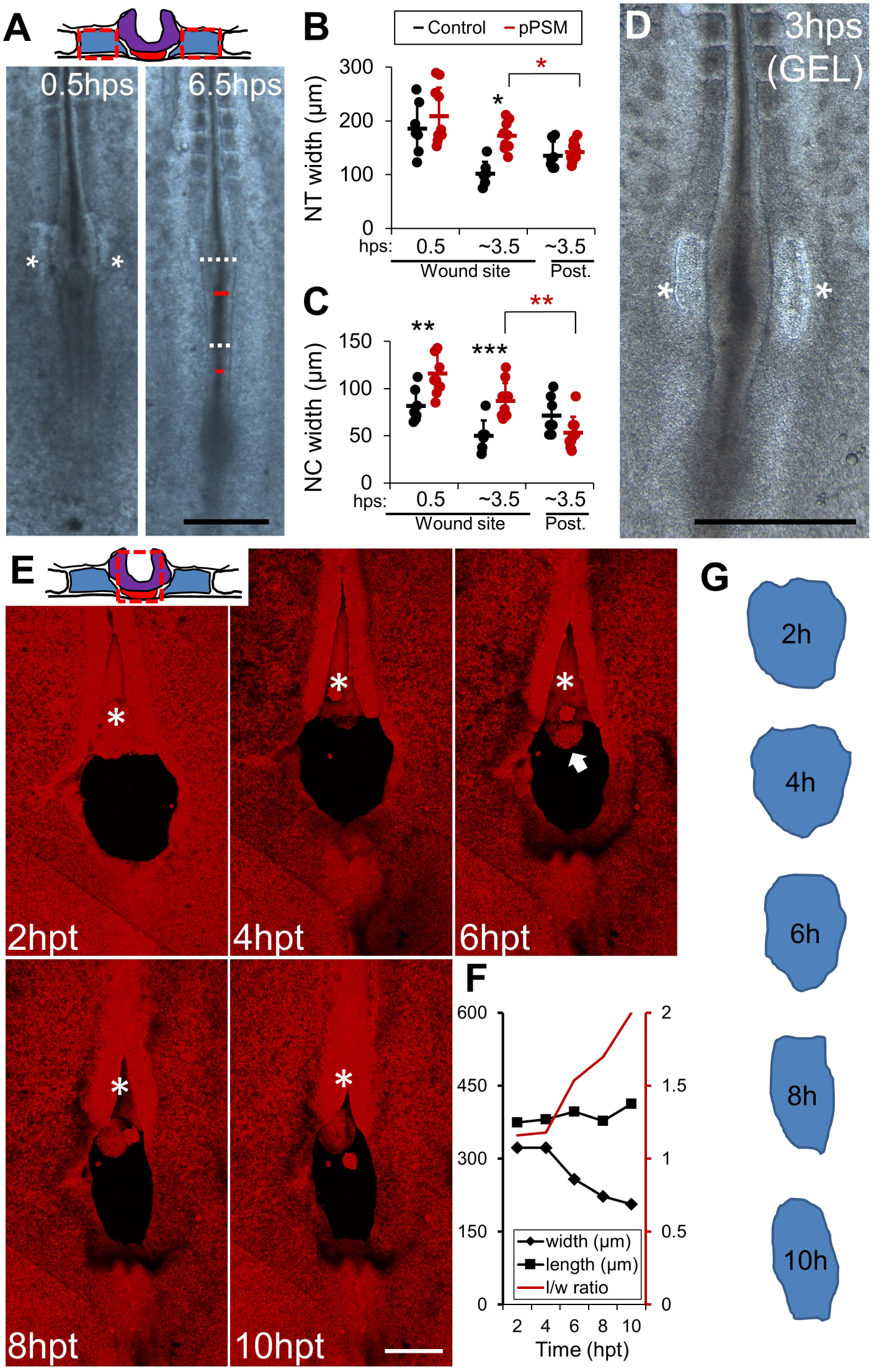
Posterior PSM exerts a lateral compression on axial tissues. (A) Ablation of the pPSM. Red dashed boxes on the cross-section schematic mark ablated tissues (same for following figures). Embryo images show axial tissue dynamics after pPSM ablation. Darkened line structure in the midline of the embryo delineates NC (red lines span the width). Lighter regions on the side of NC show the neural folds on the further dorsal side (white dashed lines span the width). Asterisks: the closing PSM ablation sites. hps, hours-post-surgery. Scale bar: 500μm. (B-C) NT and NC width shortly after pPSM ablation and later (Control n=7, pPSM n=10). Asterisks indicate significant difference (p<0.01, t-tests, unpaired (black), paired (red)). “Post.” on the horizontal axis indicates a control area measured posterior to the ablation site. (D) Gel replacement of pPSM. Asterisks, gel implant sites. Scale bar: 500μm. (E) Gel implant in place of axial tissues followed by confocal timelapse. Images are maximum projections of Tg(CAG:GFP) z-stacks (dorsal view). Asterisk shows the convergence and closure of NT anterior to the cut site. Arrow marks protruding NC underneath. Scale bar: 200μm. See also Movie S2. (F) Quantified shape change of the gel implant in (E). Length (l) is measured along the midline, width (w) is measured following the widest line perpendicular to the midline. Consistent narrowing was observed in n=3 movies. (G) Contours of the gel implants in (E).

### Cellular basis of PSM compression, axial convergence, and elongation

To investigate how pPSM creates compression on axial tissues, we constructed a 2D agent(i.e. cell)-based computational model (Figure 3A, Movie S3). The model contains a field of PSM cells subdivided by the axial tissue in the midline, and the posterior PD that supplies new PSM cells (for the feasibility of simulation the number of cells is small compared to the real tissues but roughly proportional between paraxial and axial). All cells occupy a volume in the tissue and push against each other upon close contact, and cell movement is countered by a resistance mimicking the effect of the viscous extracellular environment (Figures S3A-B). When a motility gradient is added across the field of PSM cells, the high motility region (corresponding to the pPSM) shows a diffusion-like expansion (Figure S3C) on the tissue level that reproduces observations in isolated PSM explants (Palmeirim et al., 1998). When this expansion encounters a lateral boundary, such as the axial tissues, a pressure gradient parallel to the motility gradient is recorded (Figure S3D). To test this pressure gradient *in vivo*, we replaced the aNC with gels to compare with the pNC (Figures S3E-F). In contrast to the pNC, gels replacing aNC show minimal deformations (Figure S3G). In addition, the top view area, which reflects how much the gel was compressed from the circumference over time, reduces significantly for the pNC gels while showing a slight increase in the aNC gels (Figure S3H). These data are consistent with the pPSM (but not the aPSM) generating compression in the region of maximal cell motility (Figure 1F). Our model also generates the cell density gradient observed in the PSM *in vivo* (Figures S3I-J, Bénazéraf et al., 2017), which is another consequence of the motility gradient and tissue expansion. To model axial cell behaviors, we implemented a polarity parameter that controls the shape of the axial cells. A polarized axial cell is lateral-medially elongated and intercalate with other axial cells more easily in the model, mimicking the cell shape and behavior observed *in vivo* such as neural progenitors in the NT (Colas and Schoenwolf, 2001). To model the PD, we periodically added new cells posterior to the axial end in the simulation iterations. This mimics the *in vivo* configuration where the anterior primitive streak or tailbud adds PSM progenitors at the axis end by ingression and proliferation (for simplicity, we only consider the PD placement at the posterior end of the midline, which is similar to the actual geometrical arrangement between different stages and species. In addition, PD cells behave no differently from PSM cells in our model, even though the progenitors show more complexity *in vivo* such as undergoing epithelial to mesenchymal transition [EMT, Denans et al., 2015]). A combination of the PSM motility gradient, the polarization of axial cells and the cell addition from PD produces sustained and marked axial convergence and elongation *in silico* (Figure 3B, Movie S3).

**Figure 3.**
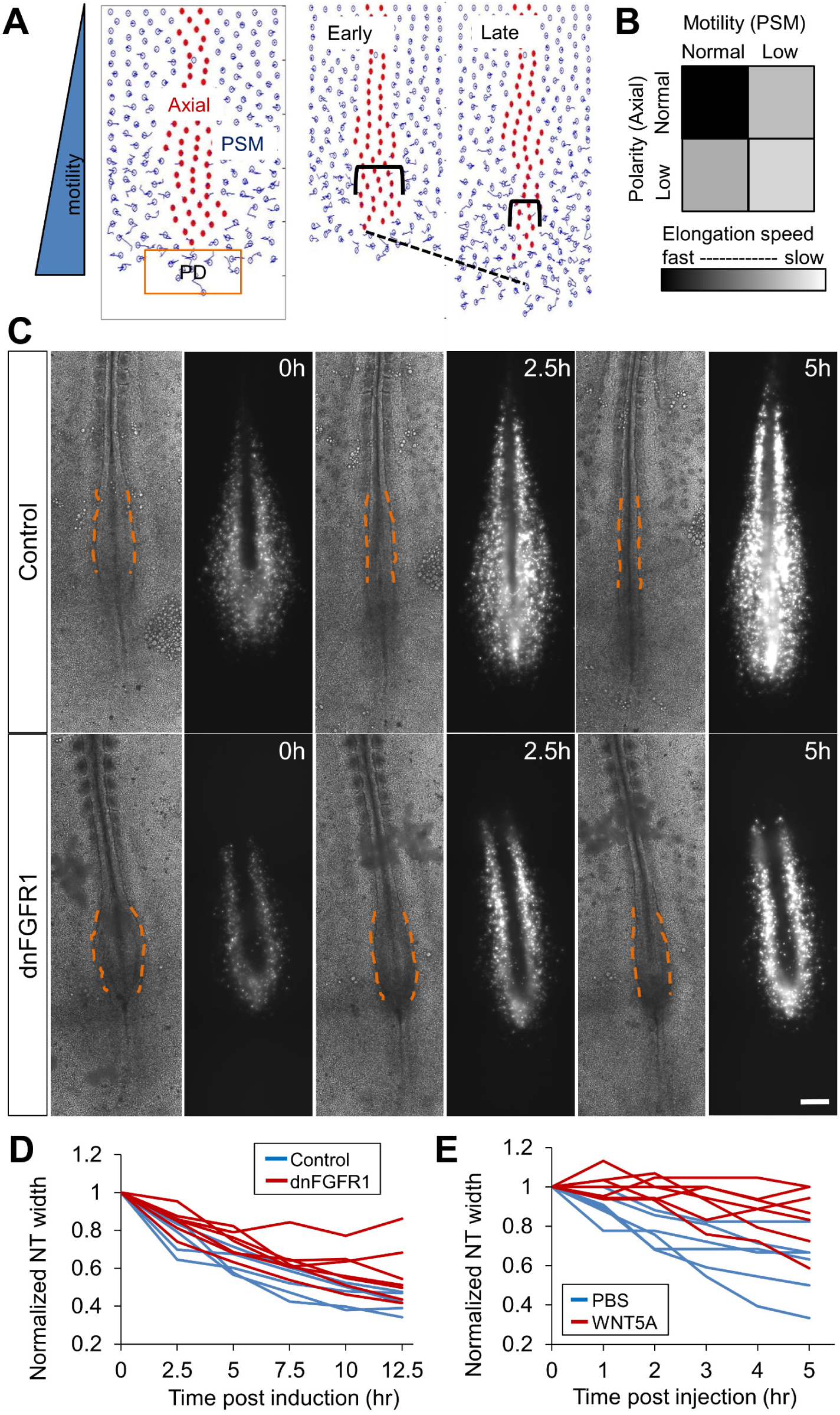
PSM cell motility and axial cell polarity contribute to axial elongation. (A) Layout of the agent-based 2D model for simulation of the PSM/Axial/PD system. NC/NT implemented as a field of cells (red) flanked by two PSMs (blue). New PSM cells are added to the PD domain (orange box) periodically. The simulation recapitulates axis elongation (dashed line) and axial convergence (brackets). See also Movie S3. (B) Elongation speed as a function of PSM cell motility and axial cell polarity in the model. Results were averaged from 10 simulation runs for each parameter pair. Each run contains 4000 time-steps. (C) Inhibition of FGF signaling by electroporating a dominant negative FGFR1 and a GFP construct into the PSM. Bright field and GFP fluorescence images of representative embryos at three different time points are shown. Dashed lines contour the narrowing pNT behind (from ventral view) the pPSM, dark strips in the fluorescent images are the NC tissue flanked by electroporated PSM cells. Scale bars: 200μm. (D-E) Convergence of pNT after dnFGFR1 induction and after WNT5A injection, respectively. Each line represents one embryo. The width is normalized to the width at the first time point. The perturbed convergence rate was significantly lower compared to control in both cases (p<0.05, t-tests).

To test the key cell behavior assumptions (motility and polarity) of the model specifically and non-invasively, we considered the molecular signals known to control these cell properties. FGF signaling is known to control the cell motility gradient in the PSM (Bénazéraf et al., 2010). We reduced cell motility by electroporating an inducible dominant-negative FGF receptor (dnFGFR1) construct and a fluorescent reporter in the PD (the anterior primitive streak, Iimura and Pourquie, 2008). The electroporation allows PSM specific expression of the transgenic construct (Figure S3K). Embryos in which FGF signaling is inhibited in the PSM exhibit a wider pNT as seen after pPSM ablation, together with slower pNT convergence and axis elongation (Figures 3C-D, Bénazéraf et al., 2010). Similar results on pNT convergence and elongation were obtained by electroporation of a dominant negative MAP kinase (dnMKK1, Figures S3L-M, Delfini et al., 2005) and by embryo treatment with the FGF inhibitor PD173074 (data not shown). These data support the hypothesis that pPSM cell movement downstream of FGF signaling collectively generates compression on the axial tissues.

pNT and pNC cells exhibit cell polarity which promotes active medio-lateral intercalation driving intrinsic tissue elongation (Keller et al., 2008; Williams et al., 2014). In amphibian embryos, interfering with PCP by disrupting the posterior WNT5 gradient reduces axial convergence and elongation (Moon et al., 1993; Wallingford and Harland, 2001). We tested whether disrupting medio-lateral cell intercalation by WNT5A also interferes with axis elongation in the chicken embryo. We injected WNT5A protein into the anterior portion of the pPSM, where endogenous expression is low (Chapman et al., 2004). This efficiently slowed down axial elongation while control injections of buffer, WNT10A at the same location, or WNT5A at the Node (where endogenous expression is high) do not have significant impacts (Figures S4A-C). Injection into the pNT at the same anterior-posterior level results in similar delay (data not shown). Consistently, the pNT in WNT5A injected embryos maintains a larger width compared to controls (Figures 3E, S4A’-B’) indicating impaired convergence. To analyze cell polarity changes following WNT5A perturbation, we electroporated a membrane tagged GFP in the NT which allows mosaic labeling of pNT cells that highlight cell shapes (Figure S5A). We also compared NT cell nuclei shapes following DAPI staining (Figure S5B). This analysis reveals that WNT5A causes a reduction of the lateral-medial polarization of NT cells in terms of cell shape (Figure S5D), while PSM cells do not exhibit cell shape polarity or change in the WNT5A condition (Figures S5C,E). Furthermore, the PSM cell motility gradient is unaffected (Figures S5F). Using live imaging and cell tracking, we found that the medial-lateral intercalation of pNT cells is greatly reduced following WNT5A injection (Figure S5G, Movie S4). These results suggest that WNT5A perturbation delays convergence and elongation by disrupting NT cell polarity. Taken together, our data show that FGF signaling in the PSM and WNT/PCP signaling in the NT provide extrinsic and intrinsic forces of axial elongation, respectively, providing support to our model assumptions and simulations (Movie S3).

### Elongating axial tissues produce a pushing force on the progenitor domain

Our perturbations show that posterior movement of the PD (which marks elongation) greatly slows down whenever axial tissues are perturbed (Figures S2B-C, S4), suggesting that the movement of the PD is not autonomous but rather requires pNT/pNC. To understand how axial elongation promotes PD movement, we resorted to our simulation. Ablating PD is predicted to slow down elongation more mildly compared to pPSM or axial ablations, which agrees with experiments (Figures 4A-B, compare to Figures 2A-C, Movie S5). We followed the average AP forces that two representative cells experience in the simulations (arrows in Figure 4A): one axial cell in the midline bordering the PD thus detecting the net force from the axial tissue and the PD, the other a paraxial PSM cell that starts off at the same AP level for comparison. The PSM cell is initially pushed posteriorly (by other pPSM cells) but then anteriorly as it transitions to become an aPSM cell, consistent with the model of pPSM expansion (Figure 4C). In contrast, the axial cell (and the PD) is continuously pushed towards the posterior. To test whether this predicted pushing force exists *in vivo*, we replaced the PD with the alginate gel. Strikingly, the gel thins antero-posteriorly especially in the middle where it is in direct contact with the axial structures (Figure 4D-F, Movie S6). This is in sharp contrast to the lateral to medial compressive deformations seen in gels that replace the pNT or pNC (Figures 2E-G, S2D-I, Movie S2), showing that elongating axial tissues indeed push on the PD. Notably, as the PD is the source that supplies new cells to the pPSM, the axial push might impact the dynamics of the PSM progenitors. In some PD gel replacement experiments, as the axial tissues advance the gel is seen to move aside from the midline and into the paraxial pPSM (data not shown), raising the possibility that PSM progenitors might respond to the axial push in a similar fashion.

**Figure 4.**
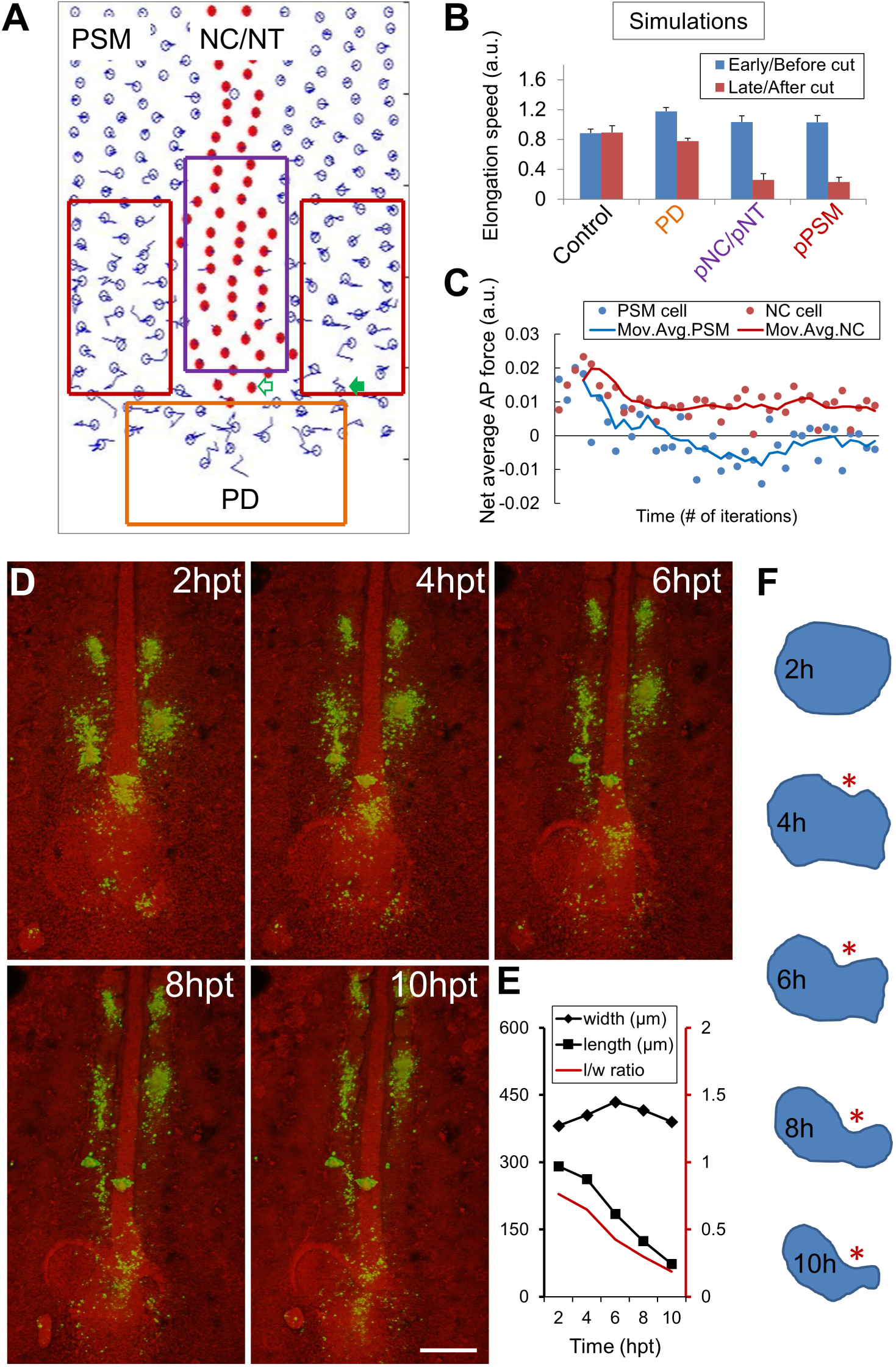
Axial tissues exert a pushing force on the PD. (A) Layout of the model showing simulated ablations (boxes) corresponding to results in (B). The green arrows indicate individual cells (empty arrow, an axial cell; filled arrow, a PSM cell) followed for the analysis in (C). See also Movie S5. (B) Axial elongation speeds (±SEM) averaged over 2000 iterations for both before and after tissue ablation. For PD, cell addition was disabled in the beginning of simulation. Each experiment includes n=40 simulations. p=0.93, control; p<0.05, other groups (paired t-tests). (C) Net forces along AP experienced by an axial cell at the anterior border of the PD and a PSM cell starting at the end of PSM. Positive indicates A to P. Data points are binned by 200 iteration windows. 4 times moving average is applied to make the curves. (D) Gel implant in place of PD followed by confocal timelapse. Green indicates spots of DiI injection to follow cell movements, spots in the pPSM can be seen expanding while the axial tissues narrow. Scale bar: 200μm. (E) Quantified shape change of the gel implant in (D). Consistent observations in n=3 embryos. (F) Contours of the gel implant in (D). Asterisk highlights the indentation where the gel is in contact with pNC.

### Axial push is required for progenitor addition to the posterior PSM

To test the effect of the axial push on the PSM progenitors in the PD, we analyzed the average PSM cell movement in the simulation. The model recapitulates the long-term trajectories of PSM cells including the medial to lateral emigration of cells from the PD into the pPSM (Figures 5A-C, see also Figure 1E). In the simulation, as there are no additional assumptions beyond cell motility and mechanical pushes between cells, the lateral emigration is promoted simply as the elongating axial tissue pushes the PD posteriorly and occupies the midline space (Movie S5). Indeed, our cell tracking (both in simulations and imaging experiments) shows that, following axial ablation, the medio-lateral (ML) speed of cells exiting the PD is reduced along with the elongation rate (Figures 5B-E, Movie S7), indicating that the rate of PSM growth is slowed down. Furthermore, the motility gradient is not sustained (Figure 5F) due to reduced addition of motile cells. These results suggest that the axial tissues not only push the PD posteriorly, but in doing so also promote cell addition into the pPSM thus extending PSM length and sustaining the motility gradient. However, a more definitive test of this hypothesis would require direct application of pushing forces on the PD in place of the axial tissues.

**Figure 5.**
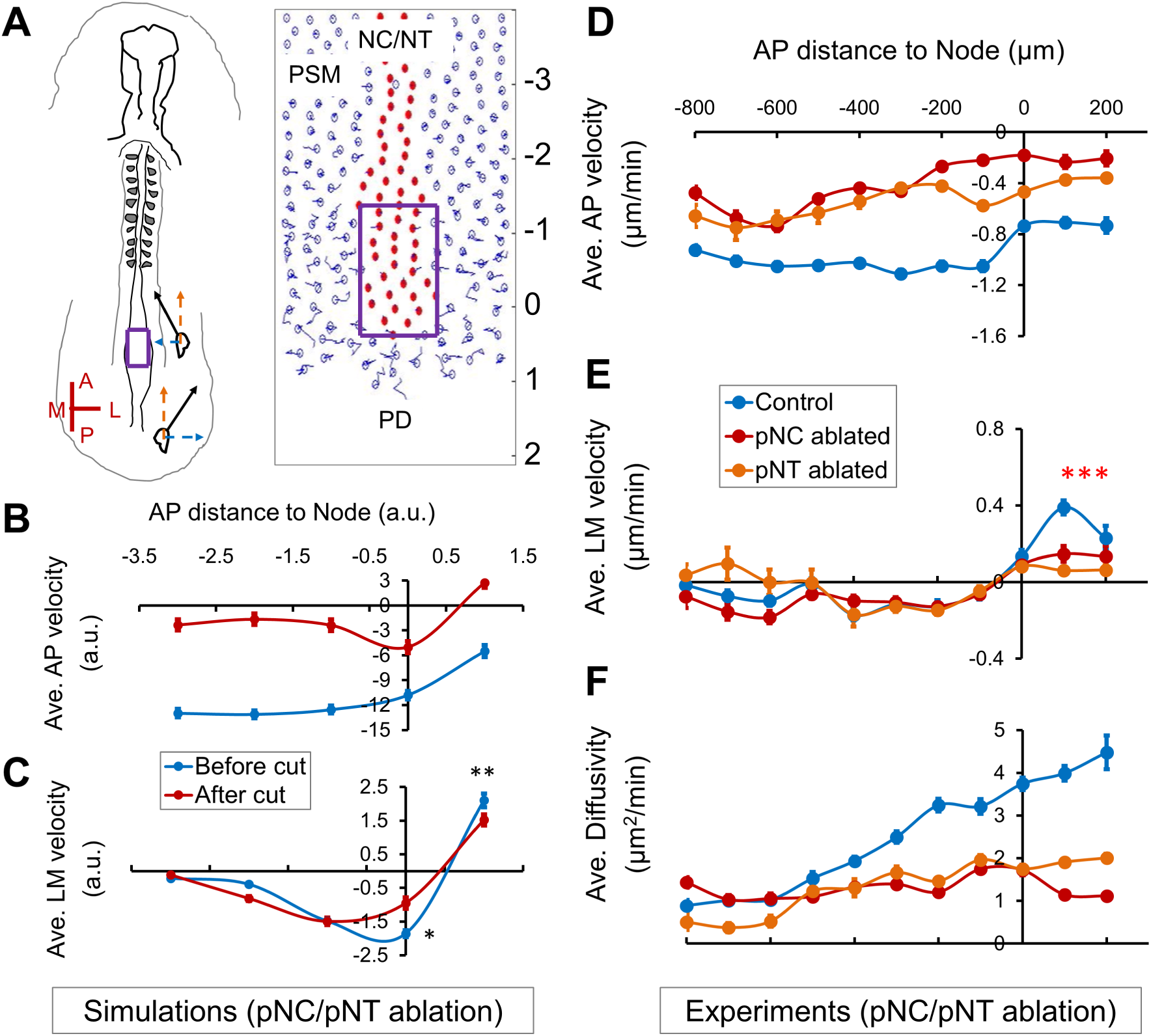
Axial push is required for cell addition from the PD to the PSM. (A) Axial ablation in the embryo and the model (purple boxes). The AP axis labels of the simulation field correspond to the labels in (B) and (C). See also Movies S5. (B) Modeled AP velocity (compare to (D)), recapitulating elongation (indicated by a negative AP speed) and slow-down of elongation after axial ablation. (C) Modeled PSM cell LM velocity (compare to (E)), recapitulating convergence, PD-exit and reduction of PD-exit after axial ablation (n=40 simulations. Asterisks, p<0.05, paired t-tests). (D-F) Average PSM cell AP (D) and LM (E) velocity and diffusivity (F) under different axial ablations. Measured and plotted similarly as in Figures 1D-F. See also Movie S7.

### Externally applied axial push rescues PD posterior movement and cell addition to the PSM

To apply an ectopic pushing force on the PD, we first performed an ablation of the pNC and then inserted a steel pin through the NT at the ablation site immediately anterior to the PD (Figure 6A). The main portion of the pin resides in the semi-liquid culture medium thus the pin can rotate around the vitelline membrane entry point (Figure 6B). In this way, the pin acts as a lever which can be controlled remotely using magnets, allowing a pushing force to be applied towards a specified direction (Figure 6C). Embryos with clean and minimally invasively inserted pins were incubated for 4 hours with or without the magnets and then imaged without the pin (live imaging was not feasible with the pin and magnet setup). In the absence of the magnet, the pin stands straight and axis elongation and PSM cell addition are slowed down as indicated by a reduction of both PD posterior movement and medial to lateral spread of labeled progenitors (Figures 6D-G). Strikingly, when the magnetic field was present to let the pin push along the posterior direction onto the PD (Figure 6C), both the posterior movement of the PD and the medial lateral spread of PSM progenitors were partially restored (Figures 6D-G). This rescue experiment shows that an external mechanical push can mimic the axial pushing force to drive the PD to move posteriorly and to add new cells to the pPSM.

**Figure 6.**
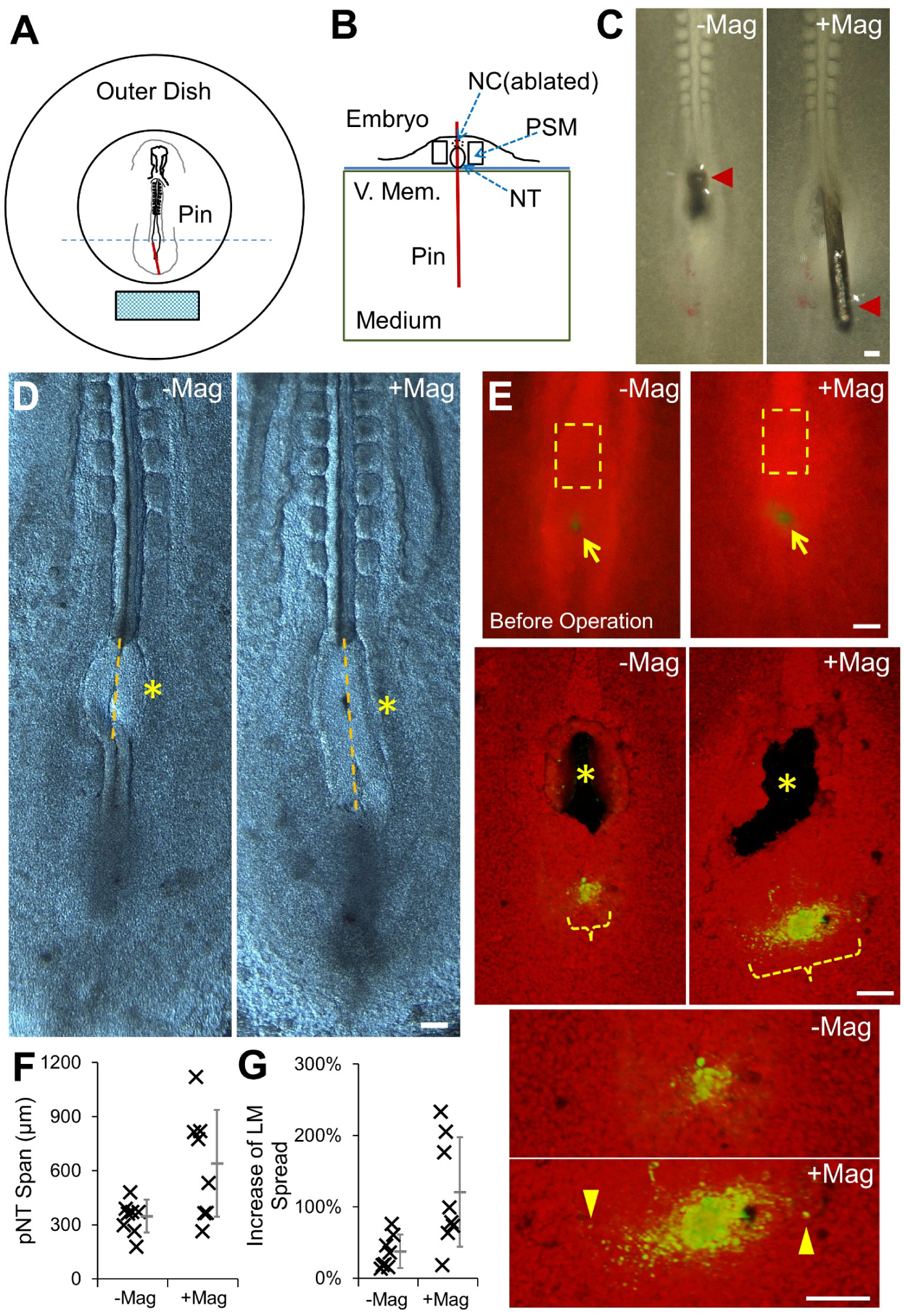
Axial push can drive PD movement and cell addition. (A-B) Top (A) and cross-sectional (B, blue dashed line in A) views of the magnetic pin set-up, respectively. Red line indicates the inserted pin and the cyan block shows the magnet (movable). The pin is perpendicularly inserted in the initial position. It rotates around the entry point of the vitelline membrane (V.Mem.). The medium is the semi-liquid albumin agar mix of the embryonic culture allowing pin rotation. (C) Example of pNC replacement with pin (left) and the push along the posterior direction after magnet is moved in range (right). Scale bars: 100μm. (D) Elongation after pNC ablation by the pin. Examples of phenotype after 4 hours with or without magnet as in (C). Pin was removed and the embryo was cleaned before imaging. Asterisks mark the pNC ablation and pin insertion site. Yellow lines mark the pNT span over the ablation site measuring PD posterior displacement. (E) Progenitor dispersion followed by pNC ablation and pin insertion. Top images show the labeled spot (DiI, green) of progenitors in the PD (arrows) occupy a narrow LM range compared to the pNC (dashed boxes) before the microsurgery and incubation. Middle images are 3D projected confocal images showing the ablation/insertion sites (asterisks) and progenitor dispersion span (Orange bracket) after 4 hours. Note that the cells have dispersed to wider LM ranges than pre-operation in both experiments. Bottom images are enlarged views showing cells that have migrated significantly laterally (arrowheads). (F-G) Quantification of elongation (F, p=0.002) and cell addition (G, p=0.011) corresponding to (D) and (E), respectively. T-tests.

### Coupling cell dynamics and tissue forces results in a self-sustaining elongation engine

Our findings show two mechanical interactions between tissues in the elongating body axis: the medio-lateral compression of axial tissues by the pPSM and the axial push on the PD by axial tissues (Figure 7A). In addition, the axial push on the PD drives new PSM cell addition to sustain pPSM growth and cell motility, thus forming a positive feedback loop (Figure 7B). This simple cross-talk ensures a tight coordination between elongation of axial and paraxial tissues, which would be difficult to achieve with separate tissue-intrinsic elongation programs. Our model predicts that co-elongation will be disrupted (i.e. length offset between tissues will emerge) as a result of breaking the feedback loop (Figure S6A, Movie S5). Indeed, ablating the PD, while itself not causing any immediate tissue length offset and allowing continued elongation (as pPSM compression remains intact in short-term), results in NC and NT that elongate further than the PSM (Figures 4D, S6B). Conversely, ablating axial tissues causes the pPSM from both sides to merge on the midline and continue to grow from the PD. This results in an axial PSM that elongates further than the NC/NT (Figures S6C-E). Our model also makes testable predictions of ablation site dynamics due to mechanical heterogeneity between tissues. For example, a square shaped opening will close preferentially along the AP axis if introduced in the PD area, or along the LM axis if introduced in the axial tissues (Figures S7A-F), similar to gel implants. Furthermore, the opening is predicted to close faster (the speed of cell density recovery after ablations) where the tissue is compressed (pNC), and slower where the tissue is expanding (pPSM), whereas for the PD that is expanding laterally and posteriorly but pushed anteriorly an intermediate closing speed is predicted (Figure S7G). Microsurgery tests *in vivo* show consistent dynamics (Figure S7H) lending further support to our model.

**Figure 7.**
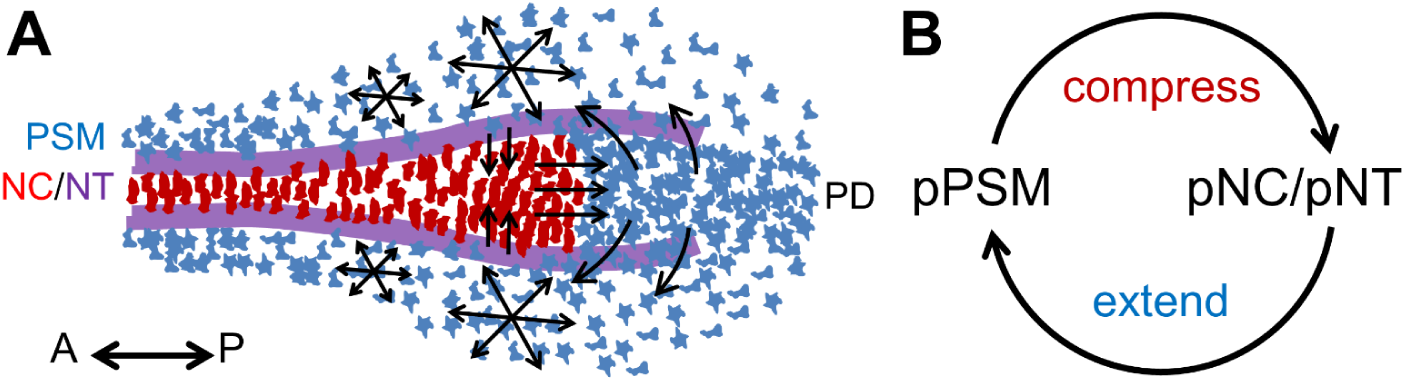
A mechanical feedback loop couples elongating tissues. (A) Schematic ventral view of observed tissue geometry and cell behavior in PSM, NC, NT and PD. Red cells: NC cells (polarized shape). Blue cells: PSM cells (mesenchymal). Purple lines: NT folds on the dorsal side. Arrows show cell movement. (B) A conceptual mechanical positive feedback loop couples co-elongation of tissues.

## DISCUSSION

### Cellular responses to extrinsic morphogenetic forces

Our study identifies inter-tissular forces as a key element in coordinated morphogenesis. This opens new avenues of investigation of specific cellular mechanisms that respond to these forces, and potential cross-talks with other cellular changes taking place at the same time. For example, the emigration of PSM progenitors from the PD to the pPSM is not only a cell movement process but also accompanied by drastic changes of the cell state. During primitive streak regression, PSM progenitors located in the epiblast in the anterior primitive streak region upregulate Wnt and FGF signaling, undergo an epithelial to mesenchymal transition (EMT) and gain enhanced motility. After tail bud formation, the PD territory becomes located at the tip of the NC and NT (the posterior wall of the chordo-neural hinge) which expresses both *Sox2* and *Brachyury* (a neural and a mesodermal marker, respectively) and continues to produce motile mesodermal precursors at a smaller spatial scale (Sun et al., 1999; Tzouanacou et al., 2009; Olivera-Martinez et al., 2012; Denans et al., 2015; Oginuma et al., 2017). In fish, the equivalent of the PD (called Dorso-Medial Zone) exhibits coherent cell movements (Lawton et al., 2013) suggesting an epithelial-like organization similar to that of the epiblast before cells migrate into the PSM. We showed that the axial pushing force on the PD regulates this lateral migration of PSM progenitors in chick. However, whether a mechanistic link exists between the axial force and the EMT of progenitors remains to be investigated. On one hand, we found that in the absence of axial tissues, the PSMs from both sides merge and continue to grow from the PD and eventually form somites (which are larger and mis-located) indicating normal PSM differentiation, suggesting that the axial force controls morphogenesis but not cell fate specification. On the other hand, we found a decrease of PSM cell motility in axial ablation experiments suggesting that the axial push may be necessary for the upregulation of FGF signaling in the PD, which is important for both EMT and PSM cell motility (Sun et al., 1999; Bénazéraf et al., 2010). Interestingly, mechanically driven FGF upregulation was recently reported in the context of branching morphogenesis in the embryonic lung explants (Nelson et al., 2017). Future studies involving transgenic markers/signaling reporters and live imaging at higher resolution in conjunction with mechanical perturbations will be important to test these possibilities.

### Force generation by cell dynamics in the posterior PSM

Recent studies have revealed the key role of the posterior PSM in the generation of elongation forces during the posterior body formation in amniotes (Bénazéraf et al., 2010; Regev et al., 2017). The physical mechanism linking the movements of PSM cells and the tissue-level expansion remains to be clarified. In our model, the force arises as highly motile cells collectively increase the space they occupy. *In vivo*, this process likely involves the remodeling of the extracellular matrix (ECM) driven by the net sum of forces exerted by single cells. Consistent with this hypothesis, fibronectin fibers in the posterior PSM follow a movement pattern that is identical to the average movement of the cells (Bénazéraf et al., 2010). Unlike the cells, the fibers do not show local random movement. The requirement of cell motility interacting with the ECM to produce tissue forces in the PSM is further supported by the observation that knocking-down fibronectins causes decoupling of PSM and NC tissues, resulting in an undulating NC and a shortened axis in zebrafish (Dray et al., 2013).

In addition to producing compression on the lateral sides, the pPSM expansion should also create an anterior to posterior pushing force alongside the axial tissues given that the underlying cell motility is random (Regev et al., 2017). The presence of this force is supported by the observation that PSM tissue can elongate on its own posteriorly without NC (Charrier et al., 1999; this study). While this pushing force appears weaker compared to the axial force according to our gel explant experiments, it might contribute to axial elongation by stretching NC/NT through inter-tissular shear forces in the posterior end. On the other hand, however, NC cells move faster posteriorly than PSM cells at most antero-posterior levels raising the possibility of NC carrying forward the PSM tissue through shear instead. To distinguish these hypotheses, it is important to design and adapt quantitative tools of mechanical measurement for very small (orders of 10-100μm) live embryonic tissues (Serwane et al., 2017).

### Robustness and self-maintenance of axis elongation in a coupled system

Our findings reveal a mechanical positive feedback loop coordinating elongation across germ layers. This situation is reminiscent of that observed during elongation of the drosophila embryo where extrinsic forces generated by the mesoderm and the invaginating gut, coupled with intrinsic forces from cell reorganization, lead to AP extension of the epithelial germ band (Butler et al., 2009; Lye et al., 2015; Collinet et al., 2015). In addition to coupling between distinct tissues during elongation, this kind of mechanical interactions may also provide robustness to other features of axis morphogenesis. For example, in our system, the FGF-dependent PSM cell motility sets a relationship between pPSM expansion and cell density, such that the compression on axial tissues increases when motile cells become more packed. This means that temporarily slower axial convergence will elicit stronger compression leading to correction. Such a mechanism could also play a role in ensuring the robustness of straight elongation (bilateral symmetry), as bending to one side will encounter increasing resistant pressure that increases with curvature, making it difficult to form large curvatures. This might provide a mechanical explanation to the recent observation that disordered cell motion in the posterior PSM is required for bilateral symmetry of paraxial mesoderm formation (Das et al., 2017).

Our model also explains the self-maintenance of axis elongation. Transplantation of the Node to an ectopic location such as the area opaca can induce a new embryonic axis which undergoes posterior elongation similar to that of the endogenous AP axis (Waddington 1932). Our model shows that elongation does not require either the formed portion of the axis or any global cues, but only a minimal set of tissues (namely pPSM, pNC/pNT, and PD) arranged around and inducible by the Node (the initial symmetry-breaking). As long as a PD (the fuel tank) that produces high motility PSM cells (combusting fuel) continues to function under the corresponding inductive signals, the positive feedback loop (the engine) can kick start locally and drive posterior Node movement, leaving a patterned body axis behind.

## ACKNOWLEDGMENTS

We thank J Chung, C. Guillot, M. Oginuma, A. Hubaud, T. Hiscock, J. Young, N. Nerurkar, S. Megason, C. Tabin, C. Cepko, M. Kirschner, P. Lenne, Pourquie and Mahadevan lab members for equipment, reagents and comments. This work is supported by a Helen Hay Whitney Foundation (HHWF) fellowship funded by Howard Hughes Medical Institute (HHMI) and a NIH pathway to independence award (1K99HD092582) to F.X. and a NIH grant R01 grant 11955884 to O.P.

## AUTHOR CONTRIBUTIONS

F.X., L.M. and O.P. planned the study. F.X. and W.M. built the models. B.B. contributed dnFGFR1 data. F.X. performed the experiments, analyzed the data and prepared the manuscript with O.P. and input from other authors. All authors participated in the interpretation of the data. The authors declare no competing financial interest.

## SUPPLEMENTAL INFORMATION

### SUPPLEMENTAL FIGURES

**Figure S1.**
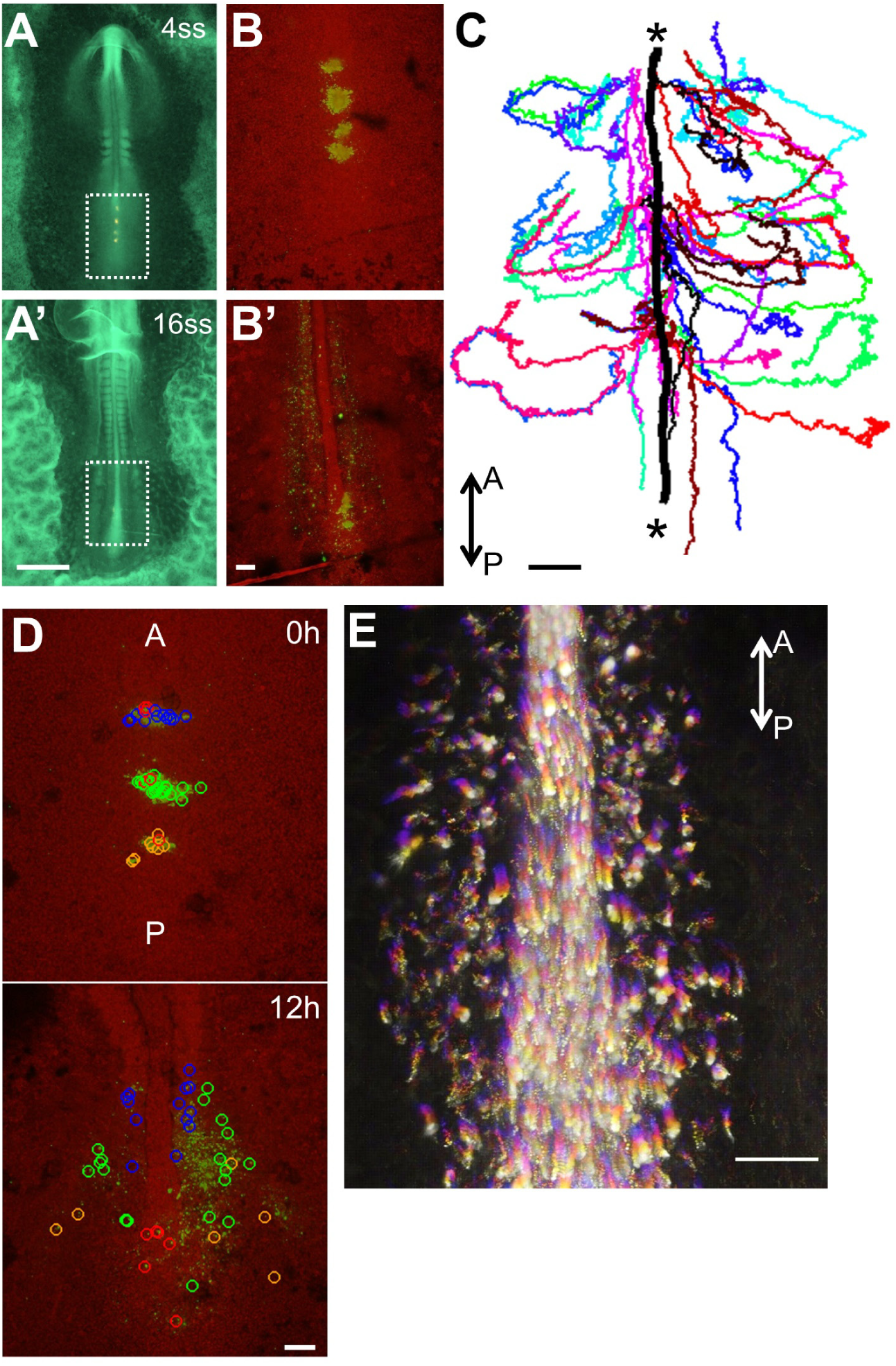
Live imaging and cell tracking in the elongating chicken embryo, related to Figure 1. (A) Whole Tg(CAG:GFP) chicken embryos (ventral view) at indicated somite stages (ss). Boxes show imaged regions in (B) Scale bar: 1mm. (B) 3D maximum projections of confocal stacks (ventral view) of the elongating axis. Ubiquitous GFP signal is shown in red, green clusters show DiI labeled sites and cells. These cells were labeled in the PD and later become PSM cells that flank the NC, which is visible in the center of (B’). Scale bar: 100μm. (C) Single cell tracks (n=46) obtained by manual tracking (Movie S1) plotted over the duration of the movie (∼15h). Asterisks mark the trajectory (bold black) of the end of NC/Node area. A, anterior, P, posterior (same below). Scale bar: 100μm. (D) Fate map of labelled cells colored as different groups posterior to NC (The blue circled cells and some green circled cells are in the PSM progenitor domain, i.e. PD, at 0h). Red circled cells either are NC progenitors or did not yet exit the midline in the duration of the movie (remaining in the PD). Somites and NC are distinguishable in the top and center of the right panel. Scale bar: 100μm. (E) Higher resolution movie (∼1h) overlay. Fire color shows progressing of time (colder is earlier). NC cells in the middle show directional movement whereas PSM cells on both sides do not show directional bias indicating short-term random motility. Scale bar: 100μm.

**Figure S2.**
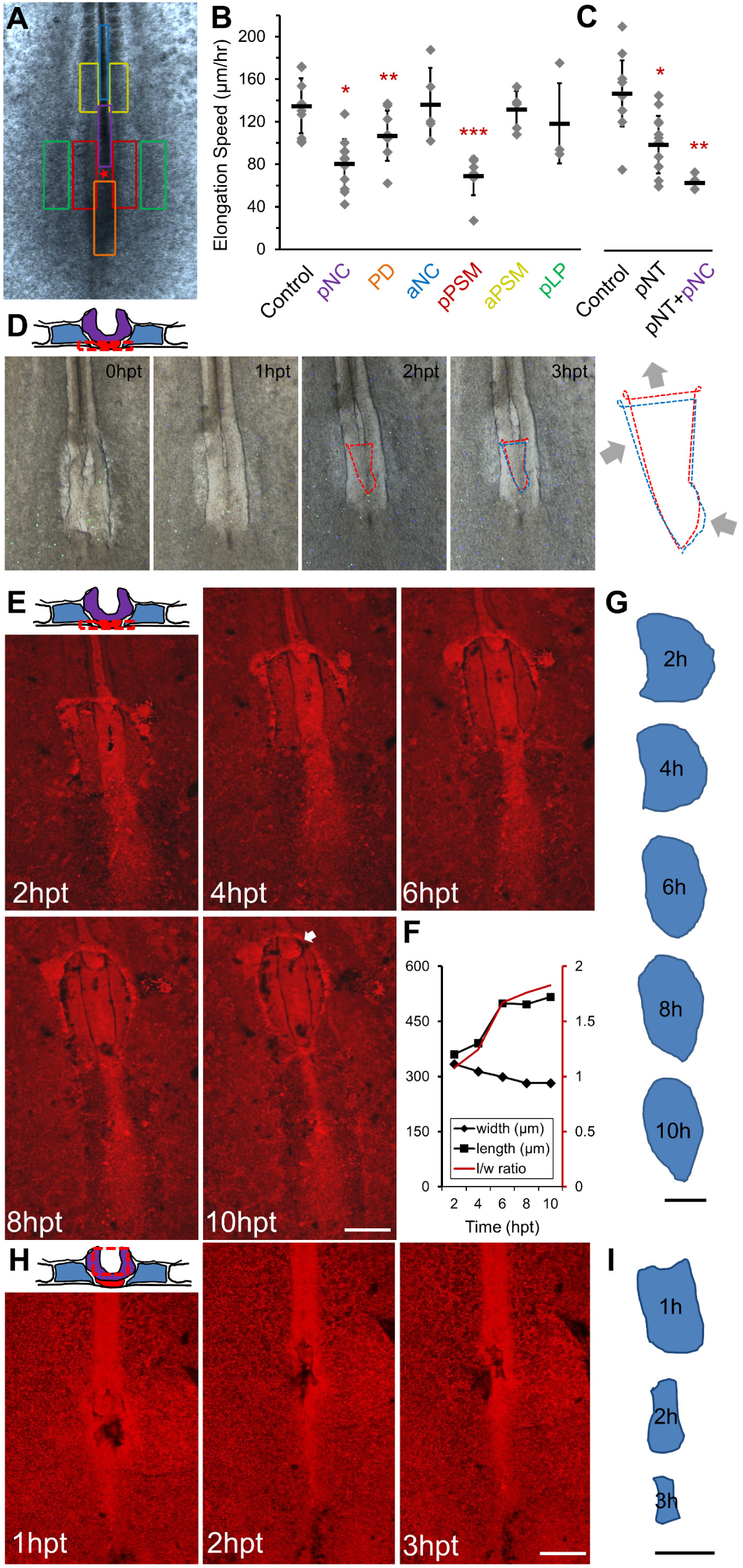
pPSM compresses axial tissues and is important for axial elongation, related to Figure 2. (A-C) Effects of tissue ablation on elongation speed. The embryo image (A) shows different ablation sites (ventral operations) with colors corresponding to elongation speed measurements in (B). (C) shows pNT ablations (dorsal operations) with its own control. Each mark represents an embryo followed over a 6-hour window after surgery. pPSM, pNC, pNT and PD ablations result in significant reduction of elongation speed (asterisks, p<0.02, t-tests). pNT/pNC double ablation results in stronger reduction than pNT alone (p=0.02, t-test, panel C). (D) Gel implant to replace pNC. Schematic image shows the operation from cross-section (ventral operations). Slight deformation of the gel can be observed through time. Dashed lines collect five fluorescent beads in the gel to a contour. The beads’ relative positions change, tracking gel deformation. The blue contour on the 3hpt image is identical to the red contour on the 2hpt image. The 3hpt contour (red) is narrower laterally than the 2hpt contour (blue), and is longer in AP (arrows). Anterior to the top. hpt, hours-post-transplant, same below. (E) Gel implant experiment in place of pNC (ventral view). Implant boundary is visible and changes over time. Only endoderm and pNC were ablated, NT is visible through the gel. Arrow marks protruding NC from anterior cut site. Scale bar: 200μm. (F) Quantified shape change of the gel implant in (E). Length (l) is measured along the midline, width (w) is measured following the widest line perpendicular to the midline. Consistent narrowing was observed in n=7 experiments. (G) Contours of the gel implant in (E). Scale bar: 200μm. (H-I) Gel implant experiment in place of pNT (dorsal view). Similar to (E), (G). Consistent narrowing was observed in n=3 experiments. pNT replacements show faster and stronger narrowing compared to pNC replacements. Scale bars: 200μm.

**Figure S3.**
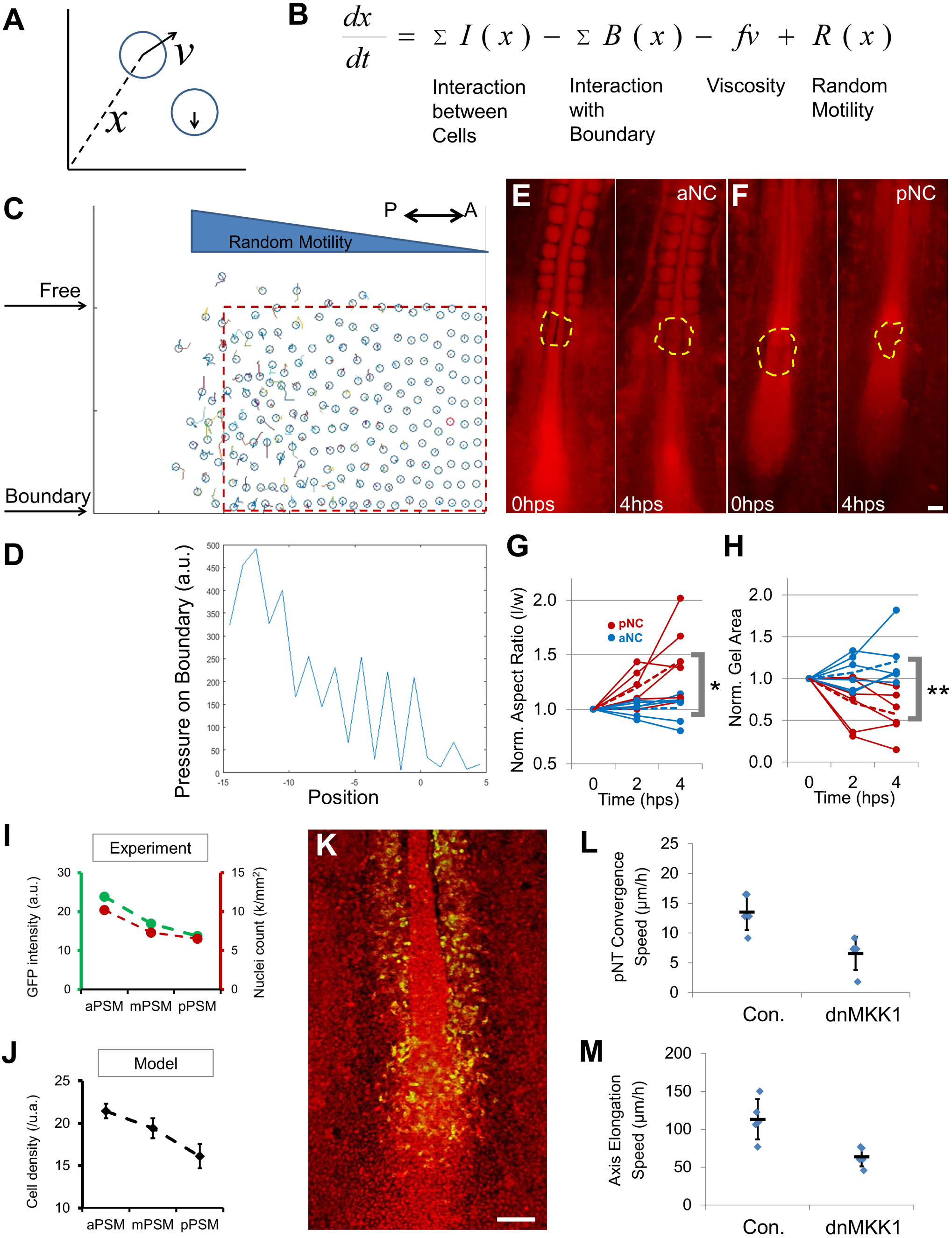
An agent(cell)-based model of co-elongation incorporating the cell motility gradient assumption, related to Figure 3. (A) Cells are defined as circles (with certain radius controlling collision volume) in a plane (tissue space, extracellular matrix). Cells have two main variable attributes: position (x) and velocity (v). Velocity is the time derivative of position. (B) For each new iteration, a velocity is generated to change the cell’s position. The velocity is determined by the forces acting on the cell, which are functions of position and current velocity. The total force is a summary of repulsive interactions between cells (a function of the cell’s position and the positions of all other cells) due to collision volume (i.e., cells have volume and cannot occupy the same area), interaction with boundary (tissue boundaries such as lateral plate), effect of viscosity (resistant force against the current velocity), and the only active movement component with a random direction and a magnitude as a function of position (e.g., the motility gradient). (C-D) A random motility gradient causes differential expansion and pressure on confining boundaries. An initial square field of cells (red box) with a gradient of random motility expands over the unconfined (free) space. On a confined boundary, a pressure gradient can be recorded. In the PSM, a similar random motility gradient is observed, and isolated PSM expands preferentially posteriorly. The pressure generated can explain the compression experienced by the pNC/pNT. (E-F) Deformation of alginate gel implants after surgical ablation of aNC and pNC, respectively. hps indicates hours-post-surgery. Fluorescent signal (red) is GFP. Yellow dashed lines measure the shape and area of the gel. Scale bar: 100μm. (G) Normalized gel aspect ratio change. “l” measures the length along AP, “w” along LM. l/w ratios for each sample was normalized by the 0 hps data point. Dashed lines are average ratios. Asterisks mark significant difference between averages at 4 hps (*p=0.018, t-test). (H) Normalized gel top view area change. Similar to (G), l*w was normalized by 0 hps data point. 4 hps difference **p=0.004, t-test. (I) Cell density measurement by quantifying GFP intensity (green) or nuclei count using Tg(PGK1:H2B-chFP) quail line (red, Huss et al., 2015) from confocal image z-stacks. a.u., arbitrary units. (J) Model predictions of cell density along the PSM. Measured areas in the model were adjusted to match measured areas from data. Error bars: ±SD. /u.a., number per unit area. (K) Electroporation specificity into PSM cells. Image shows a representative embryo electroporated at stage 5 with a membrane-Venus construct into the anterior primitive streak (PD) of a Tg(CAG:GFP) embryo. Confocal image was taken after 24 hours. Positive cells can be seen in the PD and the PSM but not in the midline NC. Scale bar: 100μm. (L-M) Converging (conv.) speed (±SD) of neural folds towards the midline (L) and elongation speed (±SD) of the axis (M) measured in 3 hours. n=5 for control and n=6 for dnMKK1, p<0.05, t-tests.

**Figure S4.**
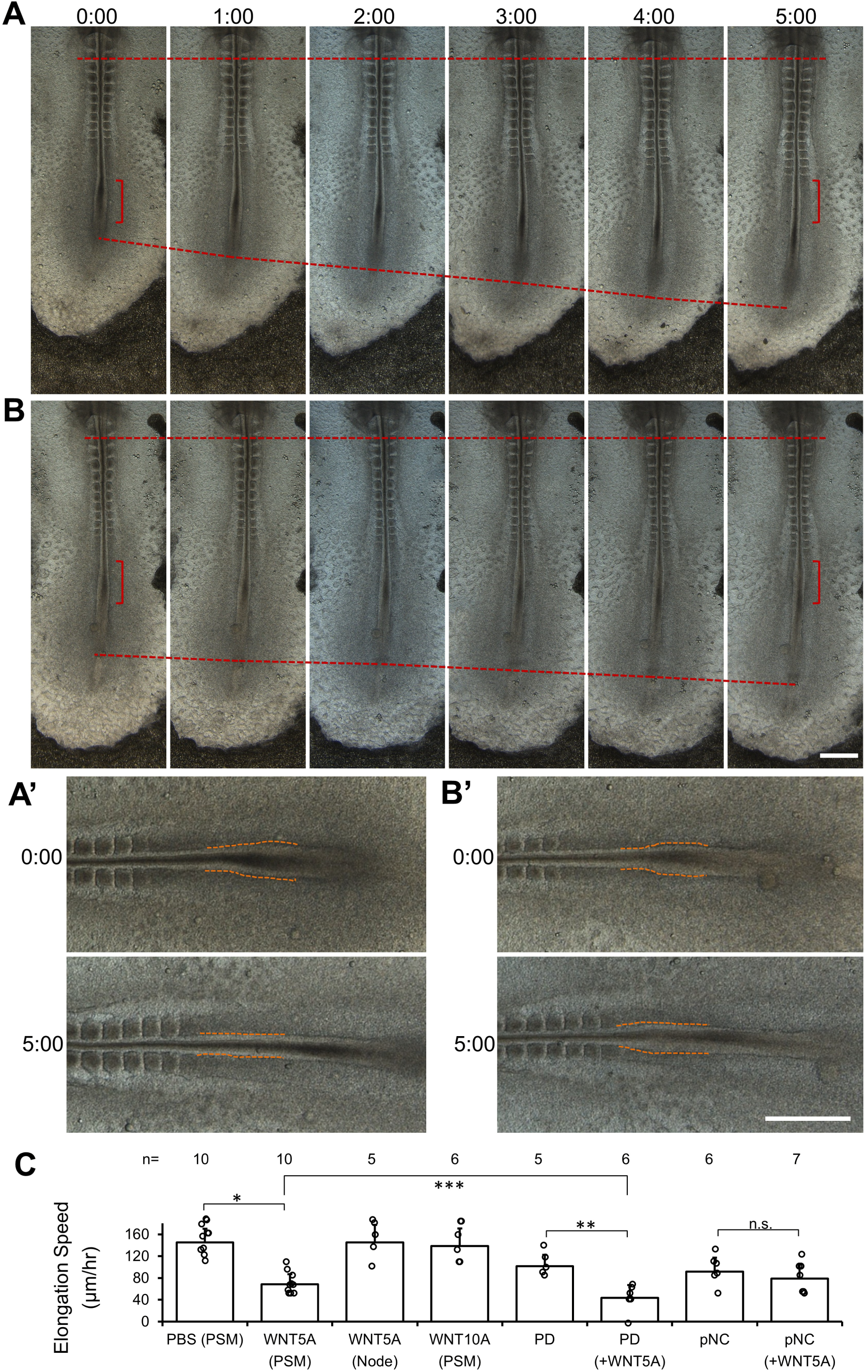
Elongation and tissue dynamics after WNT5A injection, related to Figure 3. (A-B) Time course of embryos after WNT5A injection at the PSM level (bracket). Top dashed line marks a fixed somite as the reference, bottom dashed line measures elongation. (A’-B’) are enlarged views with dashed lines showing the width of the NT. Scale bars: 500μm. (C) Elongation speeds after WNT5A injections. The first 4 columns (from left) are injections only, at the injected tissue level indicated in the parenthesis (e.g., PSM, Node). The second 4 columns couple tissue ablation (e.g., PD, pNC) with injections. Asterisks indicate significant difference (p<0.05, t-tests).

**Figure S5.**
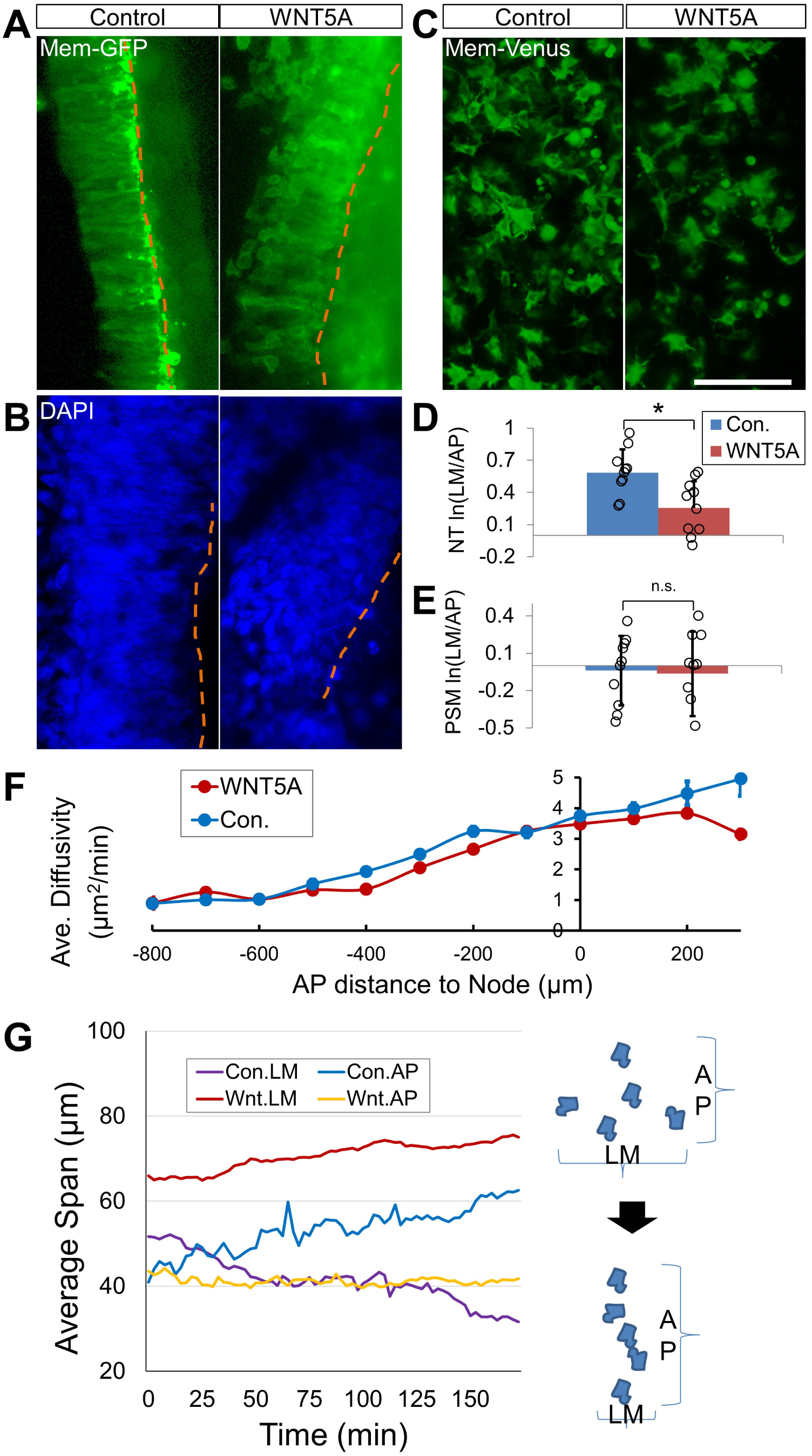
pNT cell polarity and dynamics under WNT5A perturbation, related to Figure 3. (A-C) Cell shape analysis using confocal images of cells in live embryos electroporated with a membrane-GFP construct in the NT (A, dorsal view), fixed embryos stained with DAPI in the NT (B, dorsal view), and live embryos electroporated with a membrane-Venus construct in the PSM (C, ventral view), respectively. Yellow dashed lines mark the apical/medial side of the neural fold. Scale bar: 100μm. (D-E) Quantification of cell shapes by the logarithm of the cell’s aspect ratio (lateral-medial length over anterior-posterior length, LM/AP). In (D), NT cells show LM polarization which is reduced after WNT5A injection (10 cells each were compared, *p=0.005, t-test). In (E), PSM cells show no shape polarization in either condition (10 cells each were compared, p=0.8, t-test). (F) PSM cell motility gradient in a WNT5A injected embryo. (G) Tracking of NT cell intercalation dynamics. 20 cells in each condition were tracked. Average span is calculated using the average position of all cells along either LM or AP direction. Reduction of LM span and a corresponding increase of AP span (as in illustration) indicate cell intercalation. See also Movie S4.

**Figure S6.**
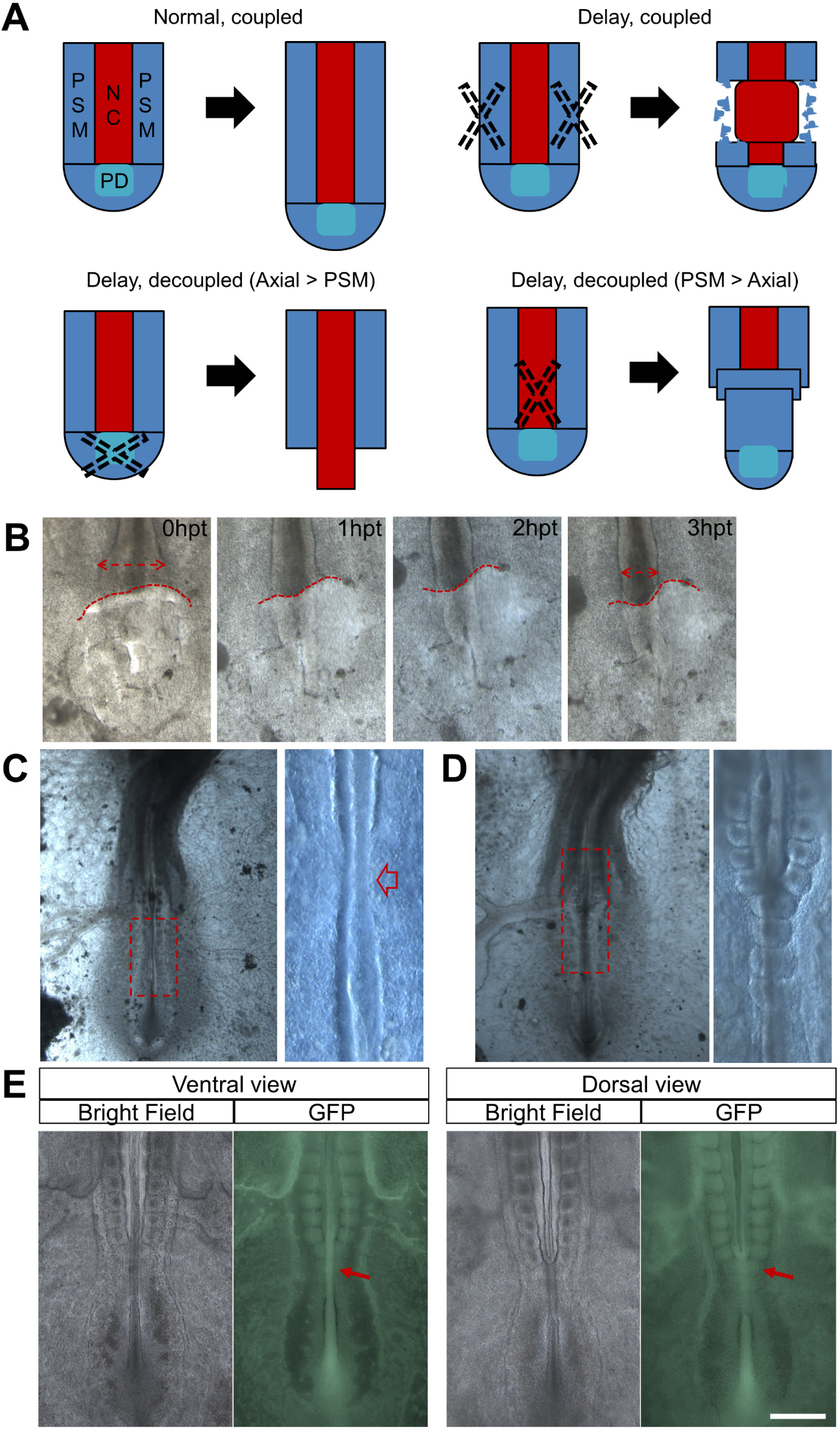
Decoupled elongation as a result of breaking the mechanical feedback loop, related to Figure 7. (A) Summary illustration of microsurgery ablations that impact elongation rate. pPSM ablation causes a local delay but does not disrupt coupling. PD ablation results in longer axial tissues and axial ablation results in longer and centered PSM. (B) Gel implant to replace the progenitor domain (PD). Double headed arrows show the narrowing of the NT. Red dashed lines show the outline of the gel in contact with pNC and pPSM. Here NC becomes longer than PSM as the gel deforms to a heart shape. (C) Left and right PSM merged at the site where pNC was ablated and elongation continued in the long term (>15 hours-post-surgery). Right panel is a zoomed in view of the red box area in the left panel. Red arrow points to the region of PSM merging. The surgery ablation of pNC usually does not completely remove all NC progenitors, thus restoration of NC further posterior is usually observed. (D) Joined PSM formed large axial somites posterior to the pNC ablation site. Right panel is a zoomed in view of the red box area in the left panel. Here PSM becomes longer than NC. Note that the NT still co-elongated with the PSM. (E) Joined PSM on the dorsal side at the pNT ablation site (arrow). Ventral view shows an undamaged NC through the ablation site. Dorsal view show joint somites forming at the NT gap. In this case PSM becomes longer than NT while NC still co-elongated with PSM. Scale bar: 500μm.

**Figure S7.**
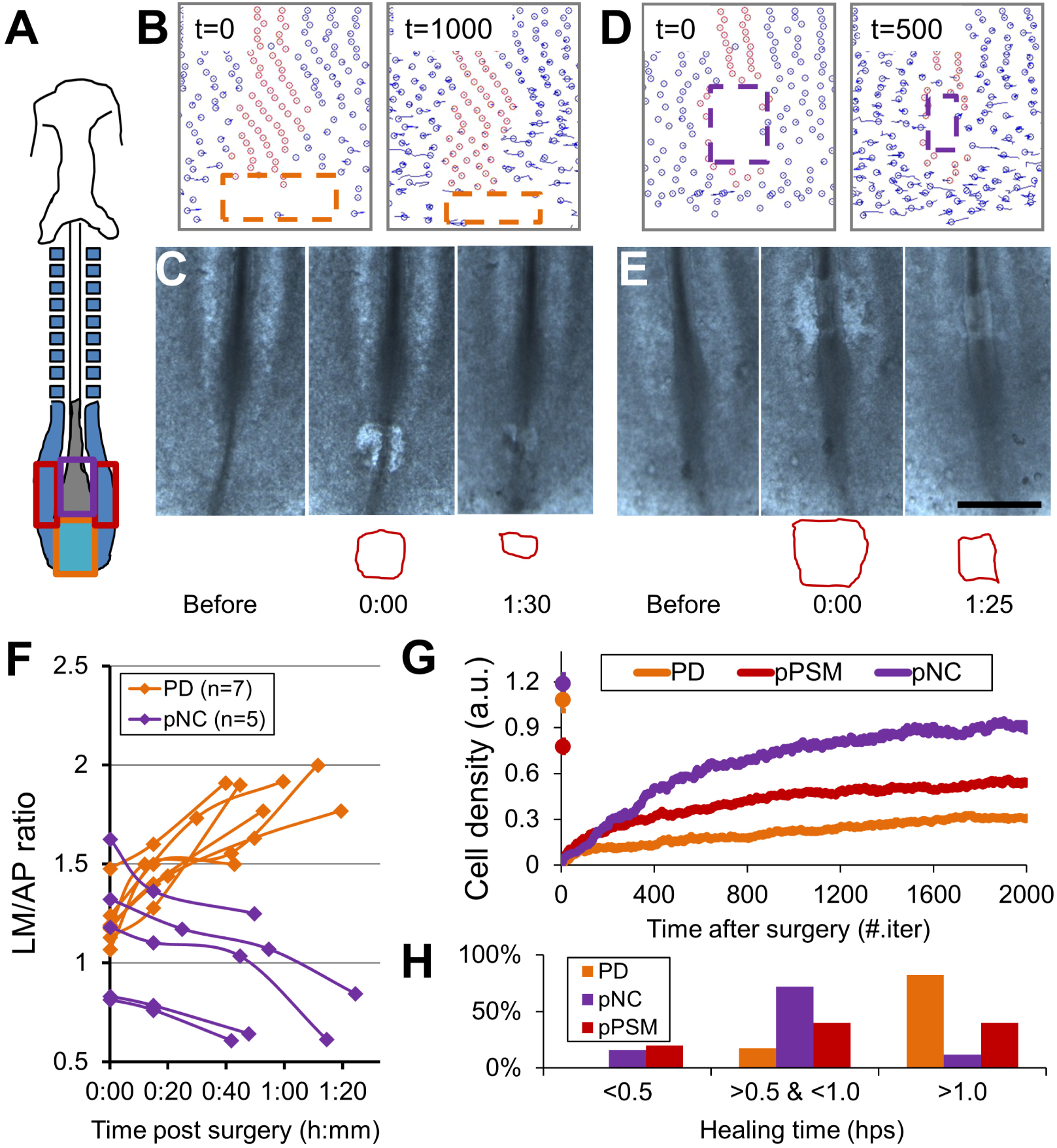
Model predictions of tissue mechanical dynamics, related to Figure 7. (A) Schematic of surgery sites on the embryo corresponding to areas in the model. (B,D) Simulations of wound closure in response to ablation of PD and pNC. Time indicates number of iterations post-ablation. (C,E) Closure of rectangle wounds introduced on embryos as shown in (B) and (D), respectively. Red contours outline the shape of the openings. Scale bar: 500μm. (F) Summary of wound aspect ratio change, LM over AP, indicating spatially differential tissue forces. (G) Simulated cell density recovery (an approximate measure of wound closure) after cell deletion in the same area. Curves show the average of 30 simulations in each case. Markers ±SD at time 0 indicate cell density before surgery. pNC area is predicted to have the fastest recovery. (H) Time of complete wound closure by ablated tissues. PD n=17, pNC n=25, pPSM n=15. hps, hours post-surgery. The majority of pNC wounds close under 1 hps, while the majority of PD wounds remain open.

### SUPPLEMENTAL MOVIE LEGENDS

#### Movie S1. Cell tracking analysis of paraxial and axial cells

The movie on the left is a 3D maximum projection (ventral view) of a Tg(CAG:GFP) (shown in red) chicken embryo imaged for the indicated duration (starting ∼3 somite stage). Green (DiI labeled) cells start as clusters at the injection site. This movie is oriented to have the anterior on the left. NC in the center can be seen moving posterior over time, followed by somite formation on both sides from the PSM. DiI labeled PSM cells disperse from the midline progenitor domain (PD) and join pPSM on both sides. On the right side the movie shows the trajectories of tracked cells of the same dataset. The solid red sphere marks the end of NC. The tracks over time are shown in Figure S1C and two time points of the movie are used for the fate map in Figure S1D. Time stamp: hh:mm. Scale bar: 100 μm.

#### Movies S2,S6,S7. Elongation and cellular dynamics under different conditions

The movies are rendered similarly as Movie S1 except anterior to the top. Microsurgical openings can be seen in the beginning of the movies of operated embryos. Detailed information for each movie is listed in the table below. Embryo stages range between 8-12ss at movie starts (number of somites). Times are hh:mm compared to movie start time. Scale bars: 100 μm.

| Movie # | Perturbation type | Perturbation time |
| --- | --- | --- |
| 2 | Axial gel replacement | -00:20 |
| 6 | PD gel replacement | -00:30 |
| 7 | none | n/a |
|  | pNC ablation | -00:25 |
|  | pNT ablation | -00:25 |

#### Movies S3,S5. Simulation of elongation under different perturbations

Movies show the 2D field (Anterior to the top) of cells (small spheres). Red cells are axial cells and Blue cells are PSM cells. In the standard simulation, new PSM cells are added at the posterior end of the axial tissues (PD). The cell addition grid is determined by one center advancing NC cell, which is allowed to move along the AP axis only to prevent bending from simulation fluctuations due to small cell number. Detailed information for each movie is listed in the table below. Times are number of iterations; all movies have 4000 iterations.

| Movie # | Perturbation type | Perturbation time | Perturbation duration |
| --- | --- | --- | --- |
| 3 | none | n/a | n/a |
|  | Reduced axial cell polarity | 1 | 1-4000 |
|  | Reduced motility gradient | 1 | 1-4000 |
|  | Reduced both motility and polarity | 1 | 1-4000 |
| 5 | none | n/a | n/a |
|  | pNC/pNT ablation | 1001 | Instant |
|  | pPSM ablation | 1001 | Instant |
|  | No Axial PD | 1 | 1-4000 |

#### Movie S4. Elongation and cell dynamics under WNT5 perturbation

The movie is 3D maximum projection (dorsal view) of Tg(CAG:GFP) (shown in green) chicken embryo after injection of PBS (control) or WNT5A. The pNT is seen in the middle undergoing convergence and elongation through NT cell intercalations. Mesenchymal pPSM cells can be seen flanking the NT. Anterior to the top. Times are hh:mm compared to movie start time. Scale bars: 100 μm.

## EXPERIMENTAL PROCEDURES

### Chicken lines and embryo preparation

Wild type chicken eggs were supplied by Charles River Laboratories. Tg(CAG-GFP) (McGrew et al., 2008) chicken eggs were provided by Clemson university (originally by University of Edinburgh). Eggs were kept in monitored 15°C fridge for storage and 37.5°C ∼60% humidity egg incubators for incubation. Whole embryo explant cultures were prepared as described (Chapman et al., 2001). All procedures have been approved by Harvard Medical Area (HMA) Standing Committee on Animals.

### Dye labeling, injections, and electroporation

3-14ss embryos were used for cell labeling. 2.5mg/ml DiI in ethanol was diluted in PBS to 0.5mg/ml before injection. The injection solution was loaded into a sharp-tipped glass needle and injected by mouth pipetting from the ventral side of the embryo into spots in different tissues (e.g., PSM, NC, the Node, and primitive streak). For over-expression of transgenes, DNA solutions were injected from the ventral side into the anterior primitive streak of HH stage 4-5 embryos, followed by electroporation using a NEPA21 type II electroporator (NEPA GENE) in PBS. The pulse setting is at 6V, 150ms interval, 25ms duration per pulse, and 3 pulses per embryo. For PSM expression, the electrode targets the anterior primitive streak (the injection site). For NT expression, the electrode targets the area slightly anterior and flanking the Node. Embryos were checked for fluorescence and development 12hr later. Poorly developed and low/miss expressed embryos were excluded from imaging and analysis. For the induction of dnFGFR1 expression in the PSM, embryos were incubated on plates with 2μg/ml of doxycyclin (Sigma). Imaging was started 2-3hrs after the incubation started to allow transgene induction. FGF signaling inhibitor PD173074 (Sigma) was solved in DMSO and diluted to 2μM in PBS before use. ∼20μL solution or DMSO control was dropped on each cultured embryo on the ventral side. For injections of PBS, WNT5A and WNT10 (R&D systems, recombinant human/mouse), 10μg/mL protein solution in PBS was loaded in a glass pipette and injected into the anterior portion of the pPSM close to both sides of the NT/NC, or into the NT apical side directly through the NC and floorplate.

### Surgeries, transplants, gel implants, and magnetic pin experiments

8-12ss embryos were used for surgery experiments. Surgeries were performed under a Leica M90 dissecting scope. Cutting was performed with an electrolysis sharpened steel or tungsten needle from either the ventral or dorsal side, for PSM/NC or NT, respectively. In the ventral surgeries, endoderm was cut open first and gently peeled aside, followed by shallow cutting into the mesoderm while avoiding damaging the neural ectoderm and epiblast. In the dorsal surgeries, the vitelline membrane over the surgery site was first slit in the middle and gently peeled on either side to make a triangular shape opening, followed by cutting around the neural plate/NT area, care was taken to minimize damage to NC and endoderm. The wound areas were gently brushed by the needle to removal cells. For transplant, the comparable region of the donor was cut intact and moved out of wound with a tungsten rod. The tissue was then transferred with a micropipette loaded with PBS to the host. A smaller opening was cut open at the desired location and the transplant was gently pushed in. For gel implant, the cleared cut opening either received an injection of 1% (w/v) alginic acid sodium (Sigma Aldrich) solution, or a small piece of pre-formed soft alginate gel (40:1 mix with 5% (w/v) calcium chloride solution). Injected alginic solution formed gel with calcium ion in the embryo culture and integrated much better than transplant of pre-formed gel. The deformation results of both methods were qualitatively similar (e.g., transplanted: Figure S2D; injected: Figure S2E) even though the transplanted gel (higher stiffness, ∼100 Pa, Banerjee et al., 2009) shows less deformation. Gels prepared with higher concentration of calcium ions or agarose gels do not show significant deformation as tracked by the movement of encapsulated beads (For example, 0.5% agarose with a modulus in the order of 1k Pa [Markert et al., 2013], data not shown). Between 20 to 45min of healing and integration was allowed before mounting the host for imaging. For steel pin (MINUCIE, stainless, No.15, ∼40μm) insertion, the tip of the pin is cut to around 1cm long. The pin is held with a tweezer and inserted between the neural folds into the culture medium from the ventral side after pNC ablation until only a small head remains level with the embryo. The embryo culture dish is placed in a bigger dish where the magnet is placed. Both distance and angle from the embryo to the magnet are adjusted by hand under a dissecting microscope to get optimal pin movement. The magnet and angle are fixed once proper pin rotation and tissue deformation under the pushing force can be seen and the configuration stays the same through a 4hr incubation.

### Imaging

Bright field and fluorescent images of the embryos were taken with a Leica M205 fluorescent microscope. Embryo culture dishes were kept in a slide box with wet paper towel. The box was kept in the egg incubator at all times except image taking (∼5 minutes for 10 embryos). For confocal timelapse imaging, the embryo (cultured as explant on filter paper) was mounted ventral or dorsal side down on a 50mm glass bottom dish (MatTek) pre-warmed to 37.5°C. A thin layer of albumin-agarose medium (200μl) was plated on the cover glass before mounting. The sample was then imaged on an inverted Zeiss confocal 780 with heating stage and environmental chamber to maintain temperature and humidity. Typical imaging set up for movies such as Movie S1: Imaging space: 1.4×1.4×0.15mm; voxel size: 1.4×1.4×2.5μm; time coverage: 15-20h; temporal resolution: 2-3min; lasers: 561nm 0.4%, 488nm 3%.

### Image Analysis

Dissecting scope images were analyzed by drawing lines over features of interest (e.g., length, wound width, notochord width, etc.) in PowerPoint (Microsoft) or Fiji (ImageJ). For elongation, the AP level of somite 2-3 were chosen as the reference point (except where it is noted differently in Figure legends). The distance increase between the reference point and the Node was used to calculate elongation speed. For confocal timelapse movies, the original.lsm (Zeiss) files were rendered into 3D maximum projection movies in FluoRender and compiled to movies in Fiji. Cells were tracked on this movie with the Manual Tracking plugin in Fiji. For cell density measurements, region of interest was selected from PSM z-stack images for fluorescent intensity or nuclei count in Fiji. Tracking results and measurements were processed in Matlab (Mathworks) with custom scripts. A track of the moving Node and a track of the anterior midline were used to linear transform the coordinates of cells to anterior-posterior / lateral-medial axes before all position and speed calculation.

### Modeling

All models were coded in Matlab. Detailed documentation is provided below. The model can be downloaded following this link: http://scholar.harvard.edu/files/fengzhu_xiong/files/fx_elongation18_modelcode.zip

#### 1. Model design and protocol

The goal of the model is to test what types of tissue behavior emerge from cells behaving under simple rules. Tissue geometry and cell activities break the homogeneity and symmetry and underlie these behaviors, although they act with a number of other interactions that should be accounted for in the model. For simplicity and feasibility, the tissue is modeled as a field of cells on a 2D plane with boundaries. The field is seeded with a finite number of cells (much fewer than in actual tissues for feasibility but geometry and proportions are kept) with user defined properties including position, velocity and cell type (Figure S3A). Time is introduced as number of iterations for the field to evolve. At each iteration, a cell receives a net “force” defined by its position, previous velocity, type, and motility (Figure S3B). This force causes it to change position and velocity. Collectively the field evolves to a new pattern. Additionally, functionalities are build-in to allow new cells to be added or specific cells to be removed during simulation, to provide tools for assessing progenitor addition and tissue surgery experiments. A chemotaxis gradient generator is also included but was kept off for the present study.

To use the model, copy all provided scripts to the working folder of Matlab. Run simulation.m. In this script, a user can define the geometrical and interaction parameters (to be discussed in detail in the next section), as well as number of iterations in one simulation, rounds of independent simulations, and whether a perturbation cut on the tissue is performed (to be discussed in detail in the section after the next). The default set of parameters produce the “Control” simulation as reported in Movie S3. To create a similar movie, specify a “moviename” string variable and use command:

~~~
“>>makemovie2(Data,N+NewN,chemoN,tipN,NCN,boundary,moviename)”
~~~

#### 2. Model assumptions

The model includes both symmetry breaking and non-symmetry breaking assumptions. The former dictates the emergent patterns of the field. The assumptions and their rationale and impact are listed below:

### Symmetry breaking

1. Geometrical layout: in the standard configuration, the model places the NT/NC cells (analogously referred to as NC cells in subsequent descriptions and the model codes) in between two flanking PSM cell groups (controlled by parameter **NCN** assigning NC cell type to the designated cell ID numbers). The progenitor domain that adds new PSM cells is placed posterior to the end of NC cells (controlled by parameter **tipN**). This layout is analogous to the actual embryo.
2. New cell addition: in the standard configuration, the model adds 80 new PSM cells to the existing field of 200 cells (controlled by parameters **N, NewN**). The cells are added periodically with parameters available to control the frequency, number and location of addition in gastrulation.m. This reflects the actual constant PSM cell production in the progenitor zone, in the current study it is either stable throughout simulation or shut down completely following removal of the progenitor domain. The interactions between these new cells to existing cells are defined exactly the same.
3. Random motility gradient across the field in the PSM cells: in the standard configuration, the model implements a simple linear motility gradient as a function of the PSM cell’s distance to the leading frontier of NC (by parameter **tipN**). The motility is produced with a random number to control direction and magnitude, with parameter **gamma** controlling overall magnitude. NC/NT cells are assumed to not have this motility. This assumption is justified by the observation of the exact type of motility gradient in the PSM in previous studies and the current study (Figure 1), and its known regulation by FGF signaling (also validated in this study, Figures 3,S3). This assumption includes cell property and a geometry asymmetry and is the **key assumption** of the model.
4. Polarity of NC cells: in the standard configuration, the model takes into account NC cell’s medial-laterally polarized shape by using a parameter **ncp** to bias the intercalation preference, implemented as differential repulsion strengths along different axes. A >1 value of **ncp** means medial-laterally elongated cell that would intercalate more easily medial-laterally than anterior-posteriorly. Simulated elongation is slower with reduced polarity and NC/NT remains wider, implying the role of cell polarity in NC elongation. This assumption is justified by the polarized NT cell shapes in previous studies and its known regulation by WNT/PCP signaling (also validated in this study, Figures S4,S5).
5. Adhesion in the anterior: an adhesion gradient from anterior to posterior in the PSM (implemented as reduced repulsion in the anterior) is used to mimic the same observed properties *in vivo*. The simulations however show that its impact on elongation is minimal. This is consistent with the observation that the main driver of elongation is the posterior PSM where adhesion is low.

### Non-symmetry breaking

1. Repulsion between cells: when cells move close to each other, a repulsion force is generated as a function of distance. This assumption shows the basic volume constrain where no two cells could occupy the same position. This assumption provides the basis for force transduction in the system and is key for implementation of several other assumptions. It alone however does not cause any symmetry break or pattern evolution. The strength of repulsion is controlled by parameter **alpha**.
2. Viscosity/friction: a resistance against the cell’s velocity is added at each simulation, to mimic the fact that cells move in a viscous tissue environment where inertia is not important. This force ensures that simulated cells do not project out of the tissues. The strength of viscosity is controlled by parameter **beta**.
3. Repulsion between cells and boundary: when cells run into a boundary it is bounced back similarly as it runs into a cell. This repulsion is not particularly important for the simulation. The parameter controlling which is set to equal **alpha**. In addition, the size of tissue field and cell field can be set by parameters **boundary** and **boundaryc**.

#### 3. Simulations

This section describes a more detailed account for the simulations. After specifying parameters described in the previous section. The user goes on to determine number of iterations (time). Sufficient time is often needed for the field to evolve into a stable state. The sum of parameters **Tcut** and **Tcut2** is the total number of iterations. This is split into two parts as the algorithm provides an opportunity of tissue cutting perturbation in between, so that the model takes the specified removed cells out of the interactions during **Tcut2**. To specify a region to cut, the user can either select one that is pre-specified in the script (cutmatrix) or write a new one. Note that this region should be a non-zero-area box (or boxes) that are within the **boundary** parameter otherwise it will not have an effect. By controlling parameter **sc**, the user can also specify how many independent runs the simulation runs.

#### 4. Result analysis

The output of simulaiton.m includes a cell array named “**Data**”, in which the cut perturbation info and the whole simulation are recorded. A user can obtain any cell’s position and transient velocity as a function of time (iteration number) and use logic index to obtain information of interest. The elongation speeds, time average cell velocities and regional density changes are computed from“**Data**”using simple scripts (not included but can be easily written).

